# XTRACT - Standardised protocols for automated tractography in the human and macaque brain

**DOI:** 10.1101/804641

**Authors:** S Warrington, KL Bryant, AA Khrapitchev, J Sallet, M Charquero-Ballester, G Douaud, S Jbabdi, RB Mars, SN Sotiropoulos

## Abstract

We present a new software package with a library of standardised tractography protocols devised for the robust automated extraction of white matter tracts both in the human and the macaque brain. Using in vivo data from the Human Connectome Project (HCP) and the UK Biobank and ex vivo data for the macaque brain datasets, we obtain white matter atlases, as well as atlases for tract endpoints on the white-grey matter boundary, for both species. We illustrate that our protocols are robust against data quality, generalisable across two species and reflect the known anatomy. We further demonstrate that they capture inter-subject variability by preserving tract lateralisation in humans and tract similarities stemming from twinship in the HCP cohort. Our results demonstrate that the presented toolbox will be useful for generating imaging-derived features in large cohorts, and in facilitating comparative neuroanatomy studies. The software, tractography protocols, and atlases are publicly released through FSL, allowing users to define their own tractography protocols in a standardised manner, further contributing to open science.

## Introduction

Diffusion tractography is a unique tool for extracting white matter (WM) pathways non-invasively and in vivo. The virtual dissection of major WM tracts enables the study of brain organisation (Catani et al., 2013; Jbabdi et al., 2015) and offers a probe to brain development (Huppi and Dubois, 2006) and WM pathology (Ciccarelli et al., 2008; Griffa et al., 2013). It further allows explorations of individual variations (Assaf et al., 2017) and cross-species variations (Mars et al., 2018b) in anatomy and connectivity. This information has functional relevance, as the pattern of *extrinsic* WM connections of each functional brain subunit to the rest of the brain are unique (Mars et al., 2018a; Passingham et al., 2002).

To be able to reliably study individual variability in WM pathways, tractography approaches often utilise protocols to extract a pre-defined set of WM tracts. Such protocols typically comprise of regions of interests (ROIs) and respective rules on using them (for instance as seed, exclusion, termination masks). They reflect prior anatomical knowledge used to guide and constrain curve propagation, reducing the chance of false positives (Catani et al., 2002; Wakana et al., 2004). Tractography protocols must be robust and reproducible, allowing reconstruction of WM tracts in a consistent manner across subjects, while respecting the underlying anatomical variation and individual differences. One approach that may be used is to define subject-specific tractography protocols (Conturo et al., 1996), considering the specific variations in individual anatomy. However, defining masks on a subject-wise basis is labour-intensive and subjective (Jones, 2008; Nucifora et al., 2012), while for large cohorts these limitations become prohibitive. The alternative to this manual approach is to define a set of standardised masks in a template space, which are then registered to the individual geometry and used in a consistent and *automated* manner for each subject.

These automated ROI-based tractography approaches have proven powerful in the extraction of a range of tracts (Catani et al., 2012; Eichert et al., 2019b; Hau et al., 2016; Hecht et al., 2015; Howells et al., 2018; Maffei et al., 2019; Makris et al., 2013, 2009; Menjot de Champfleur et al., 2013; Nowell et al., 2016; Takemura et al., 2017; Thiebaut de Schotten et al., 2011a; Zhao et al., 2016). They have further allowed for the application to large cohorts for development of tract-specific atlases (Archer et al., 2018; Chenot et al., 2019) and extraction of tract-specific features (Miller et al., 2016).

In this paper, we present a new software package with a library of standardised tractography protocols devised for the automated extraction of WM tracts, *both in the human and the non-human primate brain*. We translate prior anatomical knowledge for the human and macaque brain to a set of species-equivalent tractography rules in order to obtain homologous reconstructions for a range of WM bundles. Such equivalence can provide unique possibilities for comparative neuroanatomy and the identification of functionally equivalent cortical regions between different species (Mars et al., 2018b).

This work builds on and extends previous efforts that developed libraries of tractography protocols in the human brain (Catani and Thiebaut de Schotten, 2008; de Groot et al., 2013; Hua et al., 2008; Thiebaut de Schotten et al., 2011b; Wakana et al., 2007; Wassermann et al., 2016; Yendiki et al., 2011; Zhang et al., 2008) (see Supplementary Table 1 for a summary). A set of standard-space masks for the extraction of 20 tracts was developed by (Wakana et al., 2007). They reported high inter- and intra-rater reproducibility and suggested that some tracts may display left-right asymmetry. In (Hua et al., 2008), the authors extended this work to generate probabilistic tract atlases for 22 tracts (11 left/right) and exampled their use by investigating tract-specific abnormalities in multiple sclerosis. (Zhang et al., 2008) applied these standardised masks to 10 subjects and report high agreement between automated and manual tract segmentation approaches. Similarly, (Catani and Thiebaut de Schotten, 2008) defined standard-space masks for the reconstruction of 19 tracts (7 left/right, 5 commissural) and assessed the reproducibility of their protocols. Their work was furthered by (Thiebaut de Schotten et al., 2011b) through the extension of the tractography protocols to 31 tracts (14 left/right, 3 commissural), where good correspondence between their automated tractography technique and histological atlases was reported. (Yendiki et al., 2011) introduced a global tractography approach to reconstructing 18 tracts (8 left/right, 2 commissural) based on priors derived from an atlas. For each tract, manual tract labelling was performed in a training cohort and this provided prior information to be used in a Bayesian probabilistic framework for automated tractography. In (de Groot et al., 2013) standardised protocols for probabilistic tractography were used to reconstruct 27 tracts (12 left/right, 3 commissural) in two datasets with varying quality. More recently, (Wassermann et al., 2016) proposed a framework for describing WM anatomy and tracts which uses subject-wise anatomical segmentation, clustering and a query language to extract 57 (25 left/right, 7 commissural) tracts from whole-brain tractography and a grey matter (GM) parcellation. Their approach reduced the definition of tracts to sets of logical rules with reference to the tract position and termination, the tract route and its spatial relationship to given brain parcels.

**Table 1.**
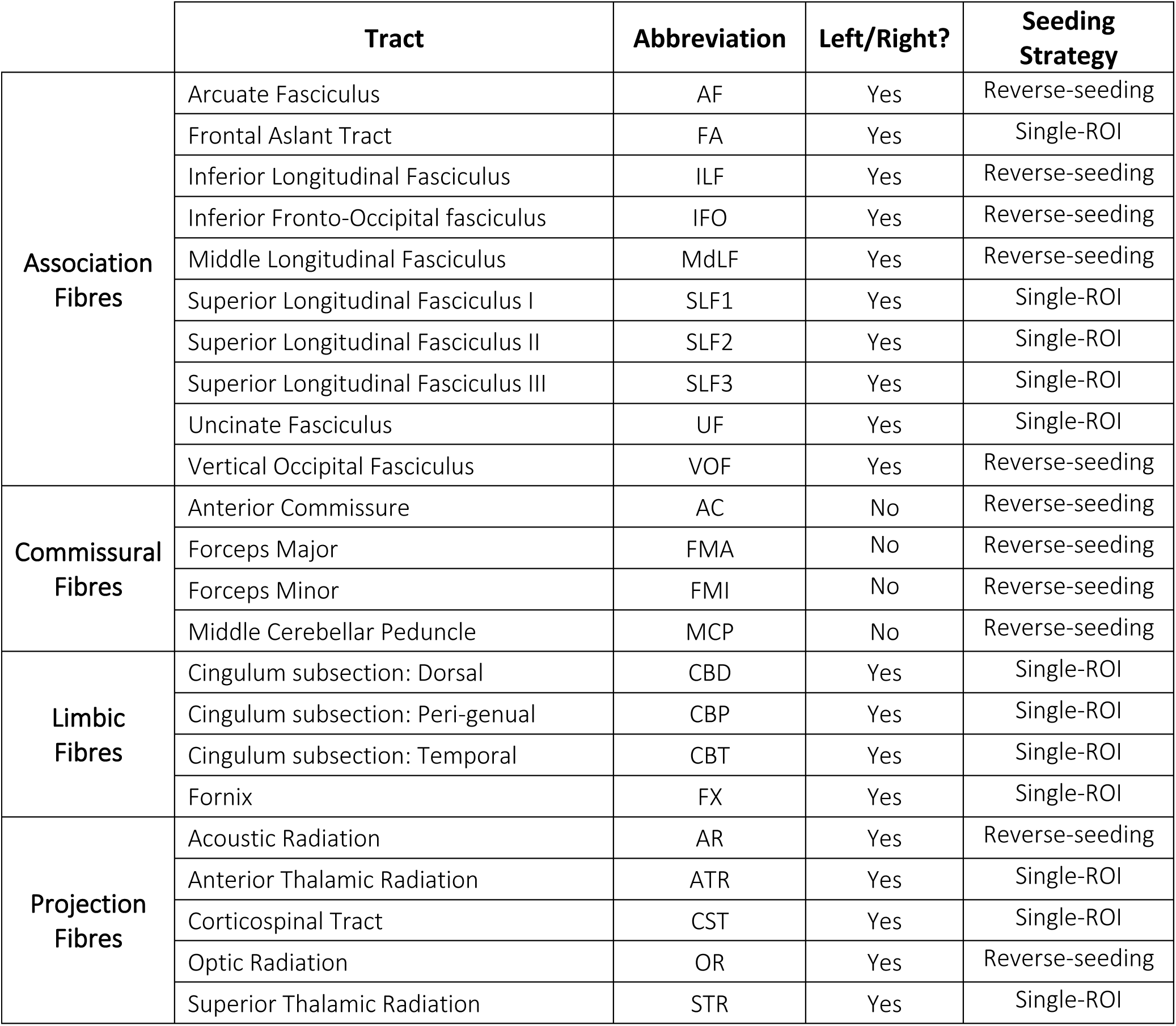
The list of reconstructed WM tracts, their correspodning tract type and the seeding strategy used.

Despite the great potential of all previous developments, none of these have targeted multiple species, which is the aim of our study. Our approach is also complementary to non-ROI-based methods for WM tract reconstruction, such as unsupervised clustering-based, e.g. (Garyfallidis et al., 2012; O’Donnell and Westin, 2007; Siless et al., 2018) or supervised methods (Wasserthal et al., 2018). The former are “data-driven”, whereas we impose prior anatomical knowledge to reduce false positives in tractography. The latter can be benefitted during training from new approaches, such as the one presented here, that aim to reconstruct anatomically known tracts in a consistent and reproducible manner.

In summary, this study devises an extended set of ROI-based tractography protocols, applicable to the human and macaque brain and also bundles these protocols in an automated tractography toolbox^1^. The contribution of the presented work is as follows: 1) we design tractography protocols for 42 tracts and we illustrate their robustness against data quality, using high-resolution data from the Human Connectome Project (HCP) (Sotiropoulos et al., 2013) and more typical data from the UK Biobank (Miller et al., 2016), 2) we illustrate generalisability of the tractography protocols to the macaque brain, 3) we derive high-quality tract atlases using these protocols both for the human brain (1000 HCP subjects) and the macaque brain (high-resolution, ex vivo datasets from 6 animals), 4) we perform indirect validation by assessing lateralisation of the extracted tracts (in humans), 5) we illustrate that, despite being template-driven, reconstructed tracts preserve individual variability as assessed via twinship analysis, and 6) we offer an open-source flexible framework for publicly exchanging tractography protocols available within FSL (Jenkinson et al., 2012). New standard space WM tractography protocols may be defined and “plugged into” the toolbox, allowing for further expansion and tract exchange, contributing to open science and reproducibility of results.

### Tractography Protocol Definition

We devised tractography protocols for 42 WM tracts (19 bilateral and 4 commissural), in a generalisable manner that allows equivalent mask definitions to apply to *both the human and the macaque brain*. The full list of tracts that are currently supported is presented in Table 1. We further packaged them into a new cross-species tractography (XTRACT) toolbox, a wrapper capable of reading the standard space tractography protocols and performing probabilistic tractography using FSL’s probtrackx2 (Behrens et al., 2007), with the option of GPU acceleration (Hernandez-Fernandez et al., 2019).

Figure 1 illustrates the main stages for a single tractography protocol. Each tract is reconstructed using a unique combination of masks, defined in standard space (MNI152 for humans and F99 for macaques). The protocols consist of hand-drawn and atlas-based masks developed and agreed upon by multiple experts (RM, KB, GD, SW, SNS). The masks include seeds (starting points of the tractography streamlines), targets/waypoints (regions through which a streamline should pass in order to be valid), exclusions (regions that serve to reject any streamline running through them) and stop/termination masks (regions that serve to stop any streamline running through them). Seeding strategies support: a) a standard single-ROI seed and b) a “reverse-seeding” approach, where a pair of seed-target masks exchange roles and the final path distributions are added. The protocol masks are transformed from standard space to the subject’s native space using a non-linear registration warp field. Tractography is performed in native space and results are directly resampled to standard space, allowing between-subject geometrical correspondence, necessary in certain contexts (e.g. atlasing).

**Figure 1.**
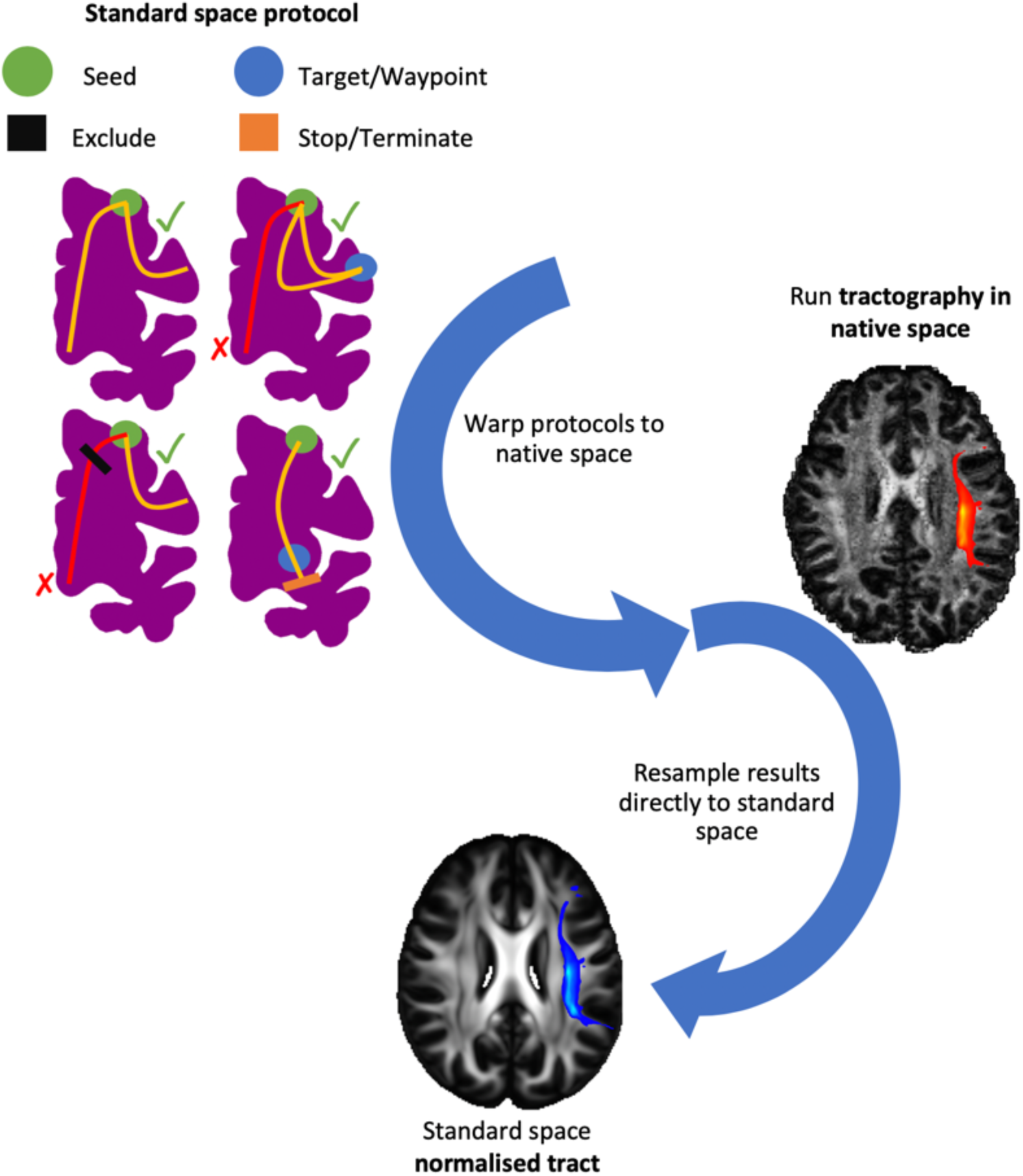
Schematic of the stages for automated tractography, as implemented in the “XTRACT” toolbox, with an example of the left arcuate fasciculus (AF) for the human brain. 1) Tractography protocol masks are defined in standard space with seed (green), exclusion (black), waypoint (blue) and termination (orange) masks (see the “Protocols” section for full details of definitions). 2) The protocol masks are warped to the subject’s native space using the subject-specific non-linear warp fields. 3) Probabilistic tractography is performed in the subject’s native space using the crossing fibre modelled diffusion data. Notice that tractography is performed in continuous coordinates in native diffusion space. Interpolation only occurs in the end when saving the result as a spatial histogram (i.e. tract visitation map). For this step nearest-neighbour interpolation is used at a user-specified spatial resolution. 4) The resultant tract stored in standard space, overlaid on the FSL_HCP1065 FA atlas.

The sections below describe in detail the protocol for each tract in consideration in the case of human tractography and any adjustments for the macaque brain. With the exception of the brainstem and commissural tracts all protocols include the midline sagittal plane as an exclusion mask to restrict fibres to the ipsilateral hemisphere.

### Association fibres

#### Superior Longitudinal Fasciculus (SLF) 1/2/3

The three branches of the superior longitudinal fasciculus are reconstructed using an extension of the approach taken by (Thiebaut de Schotten et al., 2011a). In each case a coronal plane in the region of the central sulcus within the frontal/parietal cortex is used as a seed along with two target masks. Frontally, target masks for the first, second, and third branches of the SLF were coronal sections through the superior, middle, and inferior frontal gyri, respectively, placed at the level of the posterior end of the genu of the corpus callosum. Posteriorly, a large coronal target mask in the superior parietal lobule, immediately posterior to the margin of the cingulate gyrus is used for SLF1. For SLF2 and SLF3, the second target masks are placed in the angular gyrus and supramarginal gyrus respectively. In each case, seed placement reflects the placement of the second target whilst being moved anteriorly into the region of the central sulcus. For each protocol, an axial exclusion mask was placed underneath the parietal cortex and one blocking subcortical areas prevented leaking into ventrally oriented fibres. A final coronal exclusion mask through subcortical areas posterior to the caudal end of the genu of the corpus callosum prevented leaking into ventral longitudinal tracts.

#### Arcuate Fasciculus (AF)

The arcuate fasciculus forms part of the system of dorsal longitudinal fibres, but in the human brain is distinguished by its posterior curve ventrally into the temporal cortex. The human AF was reconstructed with a seed in the supramarginal gyrus (SMG), a temporal target mask was in the WM encompassing the superior temporal gyrus (STG) and middle temporal gyrus (MTG), and an anterior target at the level of the ventral premotor cortex, posterior to the inferior frontal gyrus (IFG) and anterior to the precentral sulcus. Following the observation in the macaque that this tract runs along the fundus of the circular insular sulcus (Petrides et al., 2012) we placed a seed mask there, just posterior to the level of the central sulcus. An axial target mask was placed in the parietal-temporal WM posterior to the caudal end of the Sylvian fissure. An additional axial plane was placed in the IFG. This protocol was validated by (Eichert et al., 2019b).

#### Middle/Inferior Longitudinal Fasciculus (MdLF, ILF)

Three tracts that course along the temporal lobe were reconstructed (MdLF, ILF, IFO). The middle and inferior longitudinal tracts stay within the lateral posterior cortex. MdLF was seeded in the anterior part of the superior frontal gyrus (SFG) (Makris et al., 2009); ILF in the middle and inferior temporal gyri to account for the expansion of the temporal cortex in the human brain compared to the macaque (Latini et al., 2017; Roumazeilles et al., submitted). For the MdLF, large axial and coronal planes covering the WM in the temporo-parietal-occipital junction were used as targets, based on anatomical descriptions from (Makris et al., 2013). For ILF, a coronal plane in middle and inferior temporal gyrus is used as a target. For both protocols, exclusion masks where placed axially through the brainstem, coronally through the fornix, axially through the cingulum bundle posterior to the corpus callosum and through the entire frontal cortex. In addition, the seed mask of MdLF served as an exclusion mask for ILF and vice versa, and the ILF target mask was used as an exclusion mask in the MdLF. Additionally, for the ILF, a coronal exclusion mask was placed in the in the centrum semiovale and an axial exclusion mask covering the WM of the SMG was used.

#### Inferior Fronto-Occipital Fasciculus (IFO)

In contrast to MdLF and ILF, the inferior fronto-occipital fasciculus, also termed the extreme capsule fibre complex (Mars et al., 2016), runs more medially and courses into the frontal cortex through the extreme capsule. Extending the recipe of (Wakana et al., 2007), the seed was a coronal plane through the anterior part of the occipital cortex, the target a coronal plane through the frontal cortex anterior to the genu of the corpus callosum. An exclusion mask just behind the anterior commissure excluded all fibres except those running through the extreme capsule.

#### Uncinate Fasciculus (UF)

The bottom part of the extreme capsule contains fibres belonging to the uncinate fasciculus, curving from the inferior frontal cortex to the anterior temporal cortex. The tract was reconstructed using a seed in the STG at the first location where temporal and frontal cortex are separated, a target through the ventral part of the extreme capsule, and an exclusion mask layer between the seed and the target to force the curve. An additional coronal exclusion mask prevented accidental leaking into the fibres running longitudinally through the temporal lobe.

#### Frontal Aslant (FA)

The frontal aslant is a short tract running in the frontal lobe between the posterior part of the inferior and superior frontal gyri (Catani et al., 2012). The seed was placed sagittally in the WM of the IFG, the target axially in that of the SFG. A posterior coronal exclusion mask prevented leakage into longitudinal fibres.

#### Vertical Occipital Fasciculus (VOF)

The vertical occipital fasciculus (VOF) runs in a predominantly dorsal-ventral orientation in the occipital lobe. We used an adapted version of the recipe described by (Takemura et al., 2017). An axial seed mask was placed in the lateral part of the ventral occipital WM posterior to the anterior occipital sulcus (Petrides et al., 2012). A larger axial target mask was placed dorsally at the level of the lateral occipital sulcus. A coronal plane just posterior to the corpus callosum served as an exclusion mask to prevent leakage into anterior-posterior tracts.

### Commissural fibres

#### Middle Cerebellar Peduncle (MCP)

The middle cerebellar peduncle (MCP) was seeded in the cerebellar WM with a target in the opposite hemisphere (and their inverses). Exclusion masks were placed sagitally along the cerebellar midline and axially through the thalamus.

#### Corpus Callosum Splenium (FMA) & Genu (FMI)

We reconstructed callosal connections to the occipital lobe via the splenium of corpus callosum (forceps major, FMA) and to the frontal lobe via the genu of corpus callosum (forceps minor, FMI) using recipes based on those defined by (Wakana et al., 2007). Seed and target masks (and their inverse) for the FMA were defined as coronal sections through the occipital lobe at the posterior end of the parietal-occipital sulcus. The sagittal exclusion mask was confined to the occipital cortex and the subcortex. Additional exclusion masks though the inferior fronto-occipital WM and a coronal plane through the pons prevented leakages to longitudinal fibres. Seed and target masks (and their inverse) for the FMI were defined as coronal sections through the frontal lobe at the anterior end of the pregenual cingulate sulcus. The midsagittal exclusion mask was interrupted at the level of the anterior third of the corpus callosum and an additional coronal exclusion mask at the same level prevents posterior projections.

#### Anterior Commissure (AC)

The anterior commissure connects the temporal lobes of the two hemispheres across the midline. It was seeded in the left-right oriented fibres on the midline, with a target mask covering the WM lateral to the globus pallidae. Stop masks were placed directly underneath and lateral to the two amygdalae. A large axial exclusion mask was placed dorsal to the seed through the entire subcortex.

### Limbic fibres

#### Cingulum subsections (CBT, CBP, CBD)

Recently, (Heilbronner and Haber, 2014) proposed a segmentation of the cingulum bundle into distinct sections based on the presence of fibres connecting specific cingulate, non-cingulate frontal, and subcortical targets. We therefore created protocols for three distinct subsections of the cingulum bundle. The temporal part (CBT) was seeded in the posterior part of the temporal lobe at a section where the fibres of the cingulum are mostly oriented in the anterior-posterior direction. The target was placed posteriorly to the amygdala and stop masks were placed posteriorly and anteriorly to the seed and target masks, respectively. An exclusion mask prevented leaking into the fornix. The dorsal segment (CBD) was seeded just above the posterior part of the corpus callosum and had a target at the start of the genu of the corpus callosum. A sagittal exclusion mask in the anterior limb of the internal capsule prevented leakage into the temporal lobe. Finally, the peri-genual part of the cingulum bundle (CBP) was seeded anteriorly above the corpus callosum and a target placed below the sub-genual callosum with a stop mask placed inferior and anterior to the target. A callosal plane at the level of the rostral end of the Sylvian fissure prevented leakage into the CBD.

#### Fornix (FX)

The fornix connects the hippocampus with the mammillary bodies, the anterior thalamic nuclei, and the hypothalamus (Catani et al., 2013). The tract was reconstructed using a seed in the body of the fornix at the level of the middle of the corpus callosum and a target in the hippocampus. A callosal plane at the anterior end of the occipital cortex prevented leakage into posterior tracts and bilateral sagittal planes around the midline, at the level of the anterior tip of the thalamus prevented lateral propagation to the anterior limb of the internal capsule. To prevent leakage into the cingulum, an axial exclusion mask posterior to the splenium of the corpus callosum and a small axial exclusion covering the parahippocampal gyrus region of the cingulum are also used. We should point out that due to the relatively small size of the stria terminalis and its close proximity to the fornix, the fornix tracking may leak into the stria terminalis. This is a common issue in diffusion tractography and is yet to be overcome using approaches in line with those used in the current study (Kamali et al., 2015; Mori et al., 2017; Mori and Aggarwal, 2014; Pascalau et al., 2018).

### Projection fibres

#### Corticospinal Tract (CST)

The corticospinal, or pyramidal, tract extends from the spinal cord through the midbrain and distributes to motor cortex, premotor cortex and somatosensory cortex. The tract is seeded from the pons with a large target covering the motor, premotor and somatosensory cortices. An axial exclusion mask is used to restrict tracking to the cerebral peduncle of the midbrain. In addition, the exclusion mask includes two coronal planes, anterior and posterior to the target, to exclude tracking to the prefrontal cortex and occipital cortex respectively and a plane preventing leakage into the cerebellar peduncles.

#### Anterior and Superior Thalamic Radiations (ATR, STR)

The anterior and superior thalamic radiations connect the thalamus to the frontal lobe and pre-/post-central gyrus respectively. The anterior thalamic radiation is seeded using a coronal mask through the anterior part of the thalamus (Wakana et al., 2007) with coronal target mask at the anterior thalamic peduncle. In addition, the exclusion mask contains an axial plane covering the base of the midbrain, a coronal plane preventing leakage via the posterior thalamic peduncle and a coronal plane preventing leakage via the cingulum. A coronal stop mask covers the posterior part of the thalamus, extending from the base of the midbrain to the callosal sulcus. The superior thalamic radiation is seeded using a mask covering the whole thalamus and a target axial plane covering the superior thalamic peduncle. An axial plane is used as a stop mask ventrally to the thalamus. The exclusion mask includes two coronal planes, anterior and posterior to the target, to exclude tracking to the prefrontal cortex and occipital cortex respectively.

#### Acoustic Radiation (AR)

The acoustic radiation connects the medial geniculate nucleus (MGN) of the thalamus to the auditory cortex. It was seeded from the transverse temporal gyrus with a target covering the MGN of the thalamus. The exclusion mask consists of two coronal planes, anterior and posterior to the thalamus, and an axial plane superior to the thalamus. In addition, the exclusion mask contains the brainstem and a horizontal region covering the optic tract.

#### Optic Radiation (OR)

The optic radiation consists of fibres from the lateral geniculate nucleus (LGN) of the thalamus to the primary visual cortex. It was seeded in the LGN and the target mask consisted of a coronal plane through the anterior part of the calcarine fissure. Exclusion masks consisted of an axial block of the brainstem, a coronal block of fibres directly posterior to the LGN to select fibres that curl around dorsally, and a coronal plane anterior to the seed to prevent leaking into longitudinal fibres.

### Adjustments for the macaque brain

Although the protocols described above are such that they allow for equivalent definitions in the macaque brain, some adjustments were required to ensure anatomical accuracy. For all macaque protocols, the reverse-seeding method was used, as this was found to increase robustness in the resulting tracts. In addition, the AF and MdLF protocols were adjusted to reflect the macaque brain. In the case of the AF, a seed is placed in the caudal STG, a target directly above the principal sulcus extending posterior to 8Ad (based on the tract-tracing data of (Schmahmann and Pandya, 2006)). In addition, a target placed in the caudal STG, immediately inferior and posterior to the seed ensured tracking occurred via caudal end of the lateral fissure. For the MdLF, a single axial plane in the posterior part of the STG was used as a target.

## Materials and Methods

### Data and Preprocessing

To assess robustness across varying data quality, we utilised data from the Human Connectome Project (HCP, cutting-edge diffusion MRI) (Sotiropoulos et al., 2013; Van Essen et al., 2013) and data from the UK Biobank (Miller et al., 2016) (overall quality closer to that typically available through clinical scanners). To ensure generalisability of the protocols across species, we also utilised diffusion MRI data from the macaque brain. This data consisted of an extended set of animals used in (Eichert et al., 2019a; Mars et al., 2018b). In total, the datasets we considered consist of 1065 subjects from the HCP (all available HCP S1200 subjects that had diffusion MRI data), 1000 subjects from the UK Biobank and ex vivo high-resolution datasets from 6 macaques. For the HCP data, we removed 44 subjects with identified anatomical abnormalities from the statistical comparisons and group atlases (see the HCP quality control website for details^2^), leaving us with a total of 1021 subjects from the 1065 subjects with diffusion data.

For both the HCP and UK Biobank, we utilised the pre-processed dMRI data, available in the respective public releases (see (Glasser et al., 2013; Sotiropoulos et al., 2013) and (Alfaro-Almagro et al., 2018; Miller et al., 2016) for full descriptions respectively). Briefly, the HCP data have been acquired in a bespoke 3T Connectom Skyra (Siemens, Erlangen) with a monopolar diffusion-weighted (Stejskal-Tanner) spin-echo EPI sequence, an isotropic spatial resolution of 1.25mm, three shells (b-values=1000, 2000 and 3000 s/mm^2^) and 90 unique diffusion directions per shell, acquired twice (total scan time ∼60 minutes per subject). The UK Biobank data have been acquired in a clinical 3T Skyra (Siemens, Erlangen), consist of two shells (b-values=1000 and 2000 s/mm^2^) and 50 diffusion directions per shell, with an isotropic spatial resolution of 2 mm (total scan time ∼6 minutes per subject). In both cases, data were motion, susceptibility distortion and eddy current distortion corrected (Andersson et al., 2003; Andersson and Sotiropoulos, 2016). Nonlinear transformations to standard space (MNI152) were obtained using the respective T1-weighted images using FSL’s FNIRT (Andersson et al., 2007; Jenkinson et al., 2012), to which the distortion-corrected diffusion MRI data were also linearly registered. Concatenation of the diffusion-to-T1 and T1-to-MNI transforms allowed diffusion-to-MNI warp fields to be obtained.

For the macaque data, we combined data previously used in (Mars et al., 2018b) with newly acquired data. Data were acquired locally on a 7T magnet with an Agilent DirectDrive console (Agilent Technologies, Santa Clara, CA, USA) using a 2D diffusion-weighted spin-echo protocol with single line readout (DW-SEMS, TE/TR: 25 ms/10 s; matrix size: 128 x 128; resolution: 0.6 x 0.6 mm; number of slices: 128; slice thickness: 0.6 mm; diffusion data were acquired over the course of 53 hours). 16 non-diffusion-weighted (b = 0 s/mm^2^) and 128 diffusion-weighted (b = 4000 s/mm^2^) volumes were acquired with diffusion directions distributed over the whole sphere. The brains were soaked in PBS before scanning and placed in fomblin or fluorinert during the scan. These data will be made available via PRIME-DE (Milham et al., 2018)^3^.

Using FSL’s FNIRT (Andersson et al., 2007; Jenkinson et al., 2012), estimations of the nonlinear transformations to standard space (F99) (Van Essen, 2002) were obtained based on the fractional anisotropy (FA) maps^4^.

### Fibre orientation estimation and tractography

Prior to “XTRACT”, the crossing fibre model described in (Jbabdi et al., 2012) was applied to the diffusion data using FSL’s bedpostx and used to estimate orientations to inform tractography. This is a parametric spherical deconvolution model that accounts for the non-monoexponential decay of the dMRI signal with higher b-values. Up to three fibre orientations were estimated in each voxel along with their uncertainty. The XTRACT toolbox read the standard space tractography protocols and performed probabilistic tractography (Behrens et al., 2007). As discussed before, tractography protocols were defined for each bundle using a unique combination of seed, target, exclusion and stop masks, along with a seeding strategy (see Table 1). A number of default tractography termination criteria were also used in all protocols (curvature threshold: ±80 degrees, max streamline steps: 2000, subsidiary fibre volume threshold: 1%, randomly sampled initial fibres in case of fibre crossings in a seed location, no minimum length constraint, loop-checking and termination). A step size of 0.5 mm and 0.2 mm were used for human and macaque tractography respectively. As shown in Figure 1, the masks were warped to the subject’s native space and, after tractography, the tractography results are directly resampled to standard space. The resultant distributions are normalised with respect to the total number of valid streamlines generated (i.e. streamlines that have not been rejected by inclusion/exclusion mask criteria).

In order to obtain tract atlases, in the form of population percentage overlap, we binarised each normalised path distribution at a threshold value. The binary masks were then cohort-averaged to give the percentage of subjects for which a given tract is present at a given voxel.

### Connectivity blueprints

The estimated bundles were further used to estimate maps of “cortical termination” for each tract in consideration, using connectivity blueprints (Mars et al., 2018b). Specifically, a white-grey matter boundary (WGB) x tracts matrix **CB** was reconstructed for each subject, as illustrated in Figure 2. This was achieved by seeding from every white-grey matter boundary location and counting the number of visitations to the whole WM, giving a WGB x WM connectivity **C**_**1**_ matrix. The tracts obtained using the tractography protocols were vectorised and concatenated into a single WM x tracts **C**_**2**_ matrix. Multiplying the two matrices provides a connectivity “blueprint”, i.e. a **CB**=**C**_**1**_x**C**_**2**_ (WGB x tracts) matrix. Columns of this matrix represent the termination points of the corresponding tract on the white-grey matter boundary surface, while rows illustrate the connectivity pattern of each cortical location (i.e. how each tract contributes to the overall connectivity of each cortical location). This process was performed for the HCP subjects and the macaque datasets. The results were then cohort-averaged to produce connectivity blueprint atlases.

**Figure 2.**
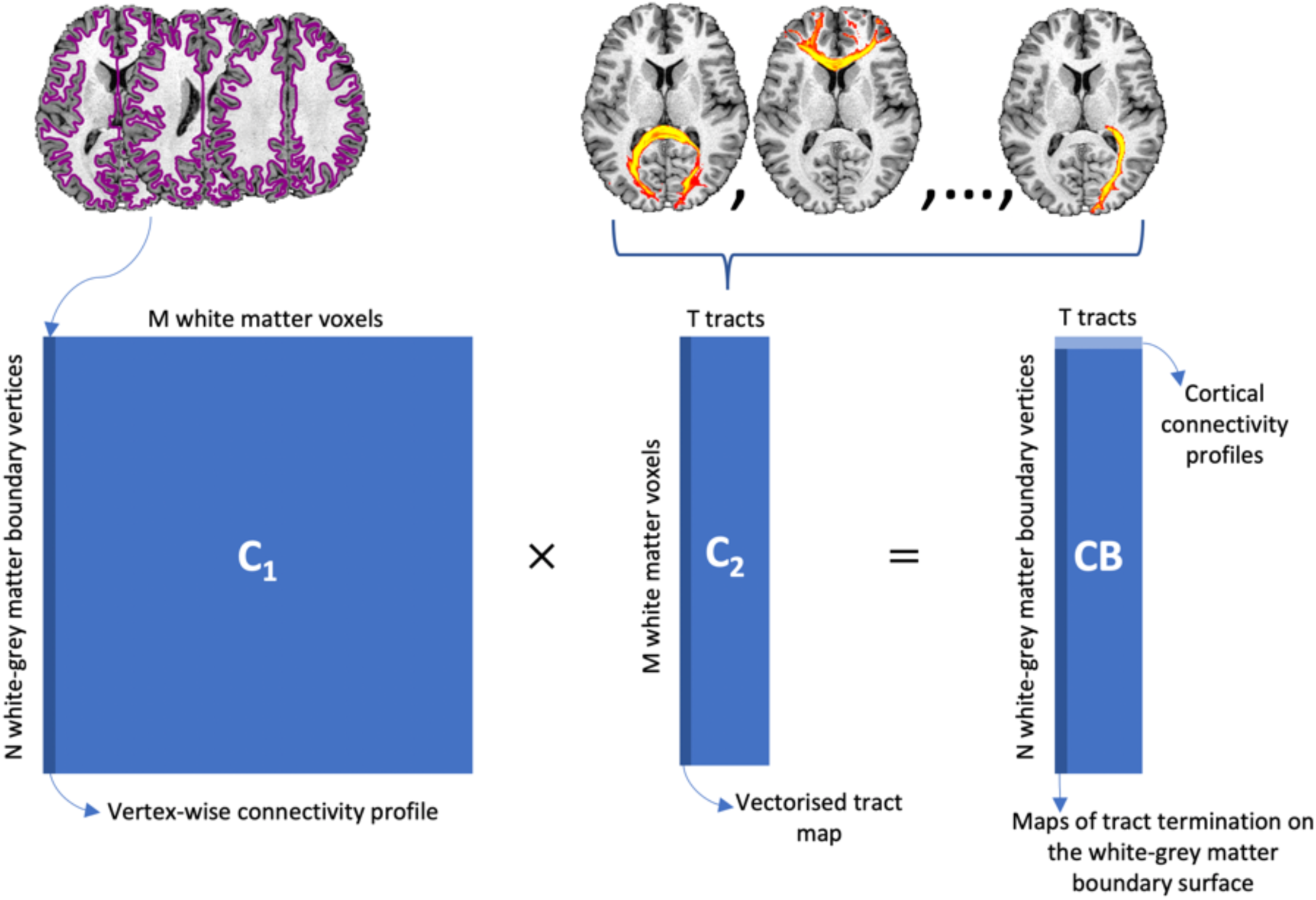
A schematic of the construction of connectivity blueprints. Tractography is seeded from the white-grey matter boundary, represented by the purple outline, and then counting the number of visitations to the whole white matter (WM), giving **C**_**1**_ (N∼60k by M∼58k). Columns of this matrix represent vertex-wise connectivity profiles. Next, the tractography reconstructions produced using XTRACT are vectorised and stacked to give a WM by tracts matrix, **C**_**2**_ (M by T=41). Multiplying the two matrices gives the connectivity blueprint, **CB** (N by T). Columns represent maps of tract termination on the white-grey matter boundary surface; rows represent white-grey matter boundary connection profiles and reflect the contribution of each tract to the connection pattern of each white-grey matter boundary vertex.

### Assessing tract lateralisation

In order to demonstrate whether our protocols produce tracts representative of the anatomical expectations, we investigate tract lateralisation using a large number of subjects. Based on the literature, it is expected that AF is left-lateralised (Eichert et al., 2019b; Nowell et al., 2016; O’Donnell et al., 2010; Panesar et al., 2018; Propper et al., 2010), IFO, MdLF and SLF3 are right-lateralised (Hau et al., 2016; Howells et al., 2018; Menjot de Champfleur et al., 2013; Thiebaut de Schotten et al., 2011a), while SLF1 is expected to be non-lateralised (Thiebaut de Schotten et al., 2011a). The literature suggests that the SLF2 is right-lateralised, however, findings are less conclusive as in some cases the reported lateralisation does not reach significance (Hecht et al., 2015; Thiebaut de Schotten et al., 2011a).

We assessed tract lateralisation using tract volume. Specifically, lateralisation (*L*) was calculated as the relative right-left volume (*V*_*r*_ and *V*_*l*_) difference, after binarising the normalised tracts at 0.5% and taking the voxel count, i.e. *L=(V*_*r*_*-V*_*l*_*)/(V*_*r*_*+V*_*l*_), in line with the literature on calculating tract lateralisation (O’Donnell et al., 2010; Propper et al., 2010; Thiebaut de Schotten et al., 2011b).

Furthermore, we explored inter-hemispheric differences in the cortical termination maps. A connectivity blueprint **CB**_**L**_, including only the left-hemisphere tracts/columns and a **CB**_**R**_, including only the right-hemisphere tracts/columns, were obtained. Both matrices were row-normalised, so that the sum of all elements in each row was equal to 1. Subsequently, we calculated the Kullback-Leibler (KL) divergence (a measure of dissimilarity) between every pair of (**CB**_**R**_, **CB**_**L**_) rows. We assessed the right-left similarity in connectivity patterns in every hemispheric location *i* using the minimum KL-divergence value obtained between all possible pairs, i.e. min(**CB**_**Ri**_, **CB**_**Lj**_), with *j* spanning all white-grey matter boundary locations.

### Respecting similarities stemming from twinship

Whilst we aimed for the automated tractography protocols to be robust against data quality, be reproducible and generalisable between species, we further tested whether they could respect features stemming from the inherent individual variability in WM anatomy across subjects. To demonstrate this, we explored the similarity of tract reconstructions within twin and non-twin sub-groups in the HCP cohort. We anticipate that monozygotic twin pairs will illustrate larger similarities than dizygotic twins and non-twin siblings, and subsequently than unrelated subject pairs, in line with the literature on the heritability of structural connections (Bohlken et al., 2014; Jansen et al., 2015; Shen et al., 2014) and as may be expected from the literature on sulcal similarities in twinship (Amiez et al., 2019). Using the 72 pairs of monozygotic twins available in the HCP cohort, 72 randomly chosen pairs of dizygotic twins, 72 randomly chosen pairs of non-twin siblings and 72 randomly chosen pairs of unrelated subjects we compared tracts across pairs to assess whether our automated protocols respect the underlying tract variability across individuals. For a given subject pair and a given tract, we correlated the normalised path distributions (in MNI space and following thresholding) of the two subjects. We repeated for each tract and the average correlation across tracts was calculated. This was then repeated for each group of subject pairs giving a distribution of average correlations for each group. We subsequently compared these distributions between the different groups.

### Respecting individual differences due to atypical anatomy

We excluded a small number of subjects from the HCP cohort-derived atlases, due to identified anatomical abnormalities, following the HCP quality control recommendations. However, we explored the performance of our tractography protocols in a number of these cases, and particularly the ability to handle atypical anatomical features and geometrical malformations. In such cases, we compared the individual tractography results with a corresponding tract atlas, anticipating that the first will respect anatomical abnormalities.

## Results

### WM tract and WGB termination atlases of the human brain

We applied the prescribed tractography protocols to ∼1000 HCP and 1000 UK Biobank subjects. Figure 3 presents the tract atlases, obtained from the HCP datasets. To obtain these atlases, the subject-specific MNI-transformed tracts were binarised and subsequently averaged to produce the population percent coverage for every tract, shown in the Figure. The atlases obtained using the UK Biobank data are shown in Supplementary Figure 1.

**Figure 3.**
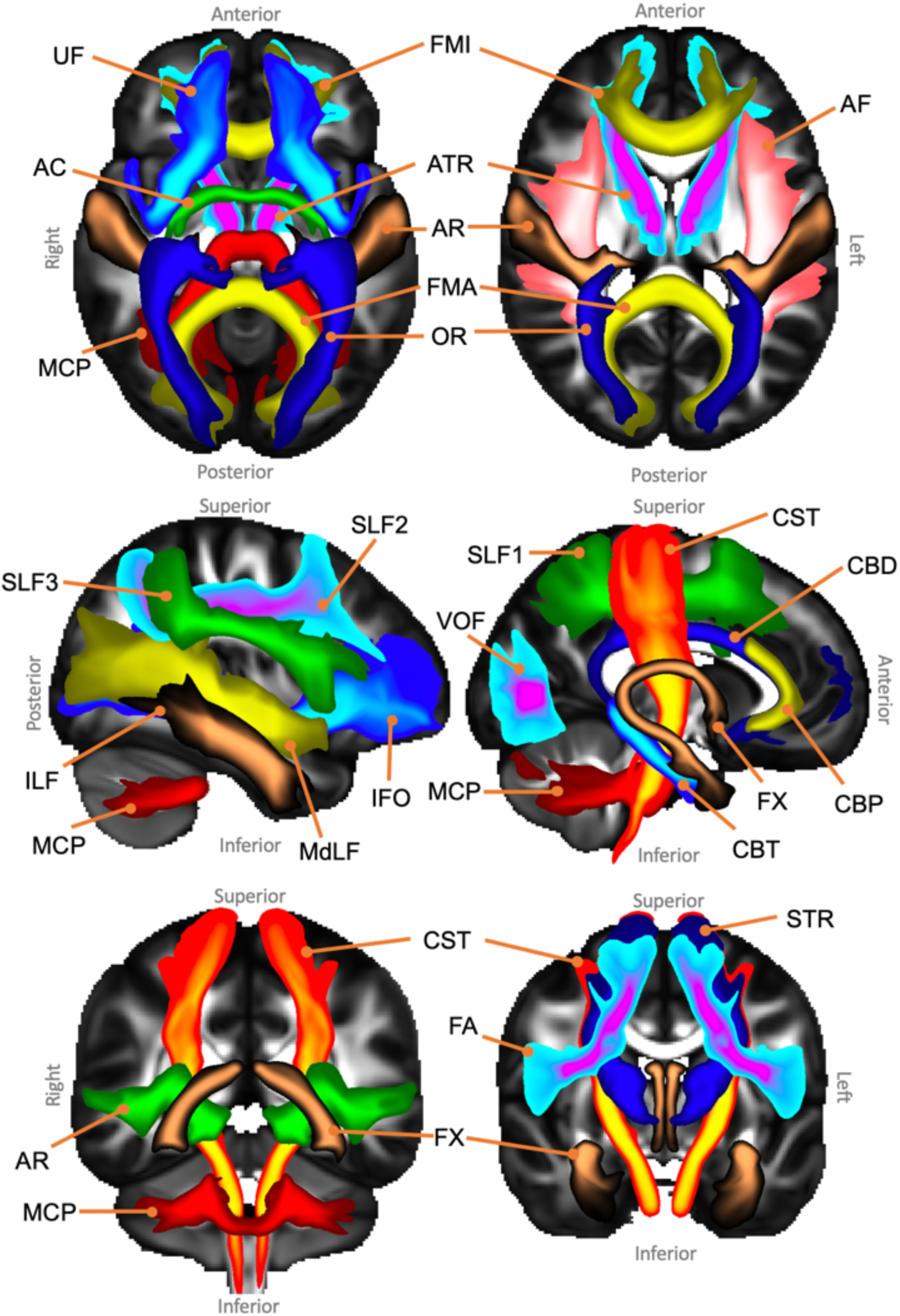
Axial, sagittal and coronal maximal intensity projections of the population percentage tract atlases (varying maximal intensity projection window lengths are applied to different tracts for visualisation purposes, display range = 5%-100% of population coverage). Association fibre bundles: Arcuate Fasciculus (AF), Frontal Aslant Tract (FA), Inferior Longitudinal Fasciculus (ILF), Inferior Fronto-Occipital Fasciculus (IFO), Middle Longitudinal Fasciculus (MdLF), Superior Longitudinal Fasciculus I, II and III (SLF), Uncinate Fasciculus (UF) and Vertical Occipital Fasciculus (VOF). Projection fibre bundles: Acoustic Radiation (AR), Anterior Thalamic Radiation (ATR), Corticospinal Tract (CST), Optic Radiation (OR) and Superior Thalamic Radiation (STR). Limbic fibre bundles: Cingulum Bundle: Peri-genual (CBP), Cingulum Bundle: Temporal (CBT), Cingulum Bundle: Dorsal (CBD) and Fornix (FX). Commissural fibre bundles: Anterior Commissure (AC), Forceps Major (FMA) and Forceps Minor (FMI). Tract atlases are created by averaging binarised (threshold of 0.5%) normalised tract density maps across subjects.

Connectivity blueprints were also derived for each HCP subject and averaged to obtain an atlas. Examples of columns of this average connectivity blueprint across all HCP subjects are shown in Figure 4, representing atlases of termination points of each tract on the white-grey matter boundary (WGB) surface.

**Figure 4.**
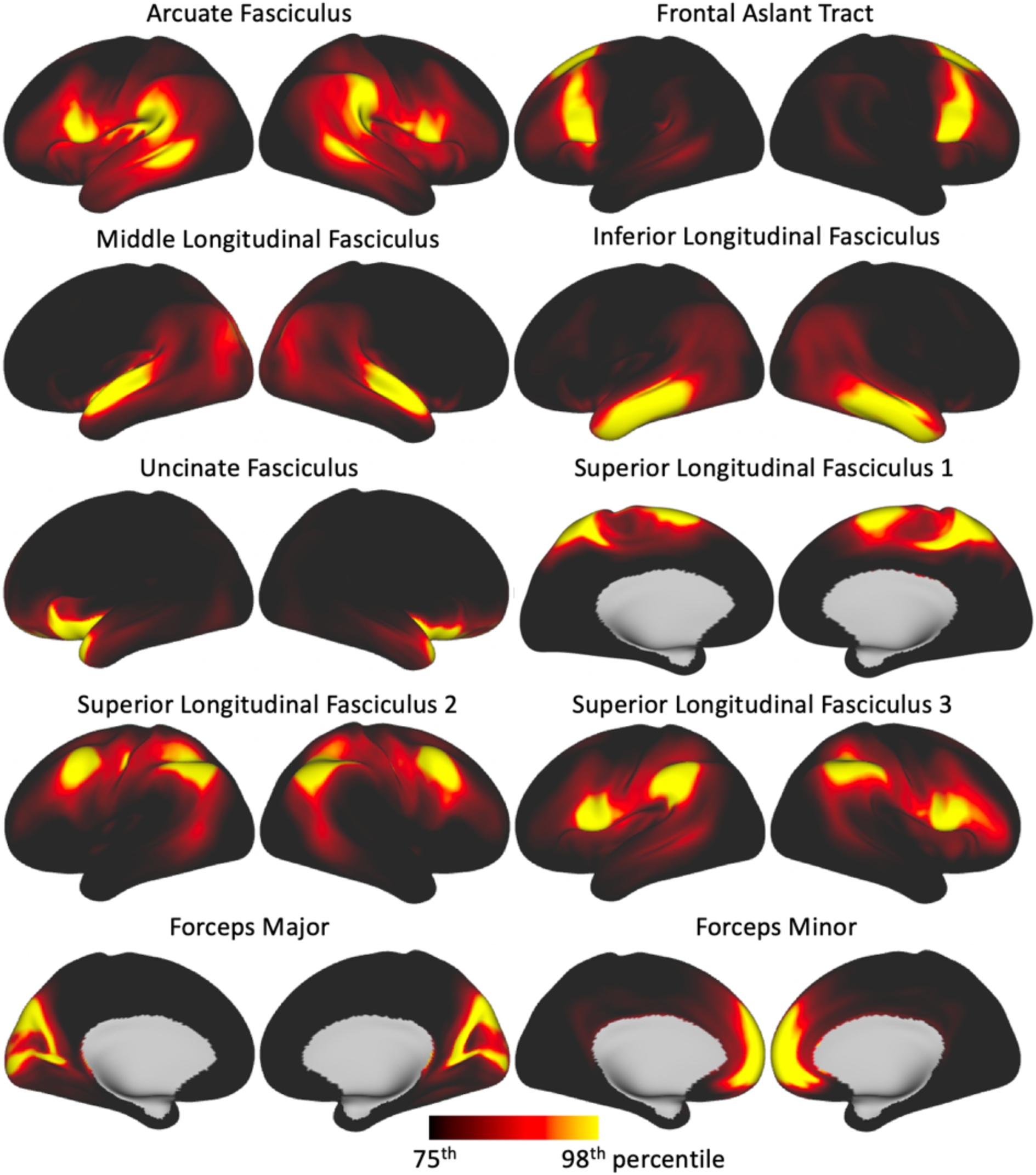
White-grey matter boundary endpoints for a subset of tracts (i.e. columns of the average connectivity blueprint) derived from the HCP cohort.

We also investigated the effect of sample size on tract atlas creation. For each of the HCP and UK Biobank cohorts, we produced tract atlases with increasing numbers of subjects and cross correlated, tract-wise, each set of atlases to a 1000-subject atlas set. Supplementary Figure 2 shows the distributions of the tract-wise correlations for each of the sample size atlases. The top plot includes an atlas set with a sample size of 10 subjects which, whilst already showing high correlations (>0.9), performs relatively poorly compared to using a sample size of 100 or greater (>0.98), i.e. the average correlation obtained from a sample size of 10 is considerably lower than that obtained from a sample size of 100 or greater.

### Robustness against datasets

To explore robustness against varying data quality, we compared tract atlases and inter-subject variability of the tract reconstructions within and across cohorts. To compare atlases, each tract from the HCP atlas set was cross correlated with its corresponding UK Biobank tract atlas (population threshold of 30% applied to each tract atlas). The average correlation across tracts was 0.80 (standard deviation = 0.07).

Inter-subject correlations were obtained by cross correlating random subject pairs tract-wise (i.e. correlating the normalised path distributions in MNI space for each tract) and averaging the correlation across tracts for each subject pair. This was repeated for many subject pairs within and across cohorts. To avoid possible family structure-induced bias in the HCP, we restricted our subjects to the 339 unrelated subjects. We matched the number of subjects in the UK Biobank data by randomly selecting 339 (gender matched) subjects. The across-cohort comparison was made by correlating a random subject in the unrelated HCP subject pool with a random UK Biobank subject, giving a distribution of 339 correlations per tract. Within-cohort comparisons gave average correlation values of 0.51 (standard deviation = 0.03) and 0.54 (standard deviation = 0.03) for the HCP and UK Biobank respectively (*p* = 8×10^-46) (Figure 5). Across-cohort comparison gave an average correlation of 0.41 (standard deviation = 0.03) which is lower than within-cohort comparisons (*p* = 1×10^-110 and 2×10^-110), yet it is comparable enough, particularly given the differences in data quality and the age difference of subjects in the two cohorts (HCP:22-35 years old, UK Biobank: 40-69 years old, mean age for our chosen subjects in HCP = 28.6 (standard deviation = 3.7) and UK Biobank = 62.6 (standard deviation = 7.5)). Indeed, we have found that tract volumes are, on average, larger in the HCP cohort compared to the UK Biobank cohort (6.5% ± 11.3% larger in the HCP cohort), which is in line with the literature on age-related changes in WM volume (Lebel et al., 2012; Rathee et al., 2016; Westlye et al., 2010). Supplementary Figure 3 shows the distributions of tract volumes for each tract reconstructed for the HCP and UK Biobank cohorts.

**Figure 5.**
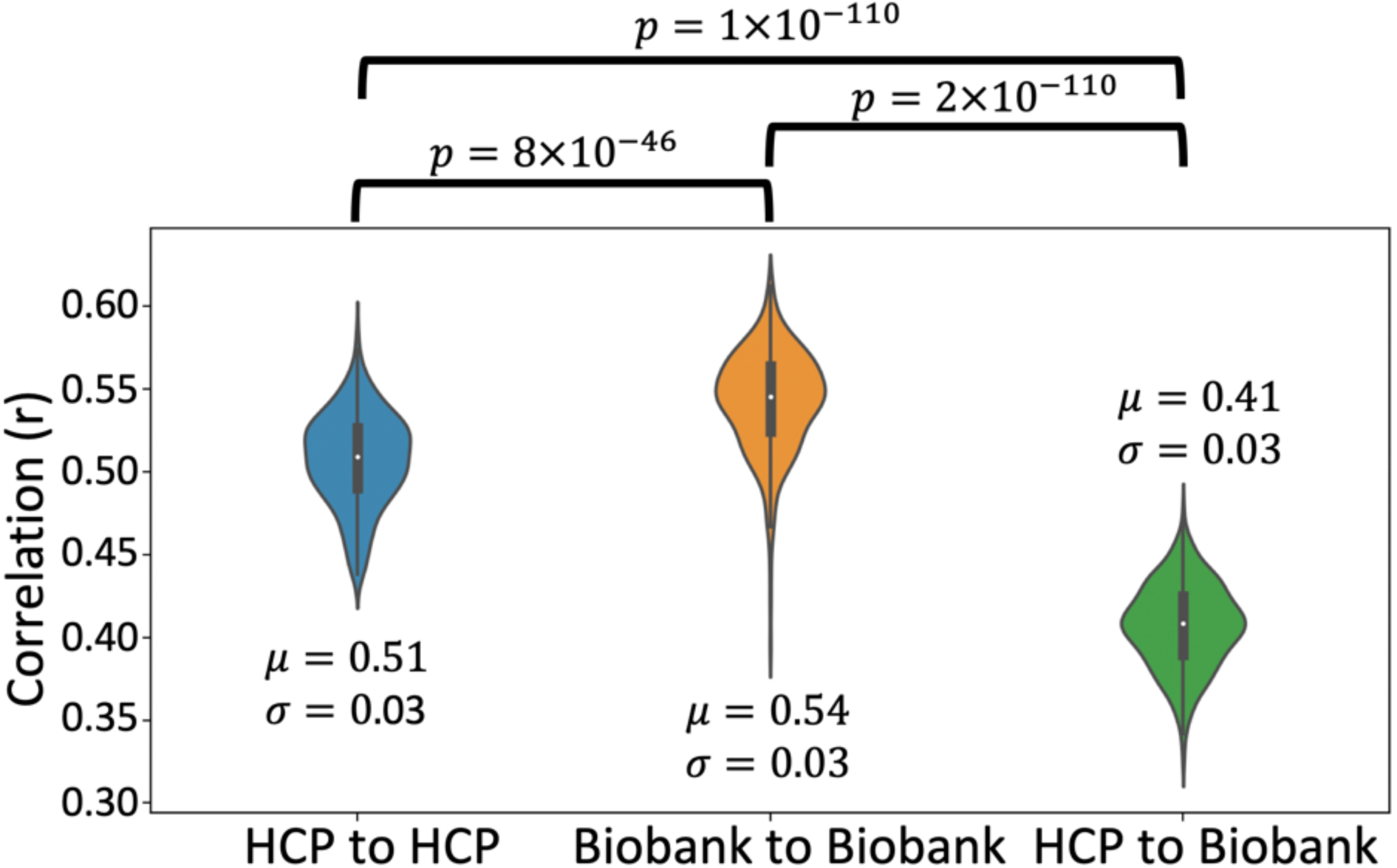
Summary of inter-cohort robustness. Plots of the average tract-wise correlations between 339 subject pairs within and across cohorts. Correlations are performed on normalised tract density maps with a threshold of 0.5%. μ is the median of the correlations across tracts and subject pairs and σ is the standard deviation. Significance is obtained via Mann-Whitney U test. Corrected p-value is 0.05/3 = 0.017.

### Generalisation across species - WM Tract and WGB termination atlases of the macaque brain

We repeated tractography as described above and obtained atlases from the macaque cohort. Our tractography protocol definitions are such that they allow the extraction of homologous tracts in both the human brain and macaque brain.

Tract atlases in the form of population percent were generated and are shown in Figure 6 and Supplementary Figure 4, while Figure 7 shows the averages of the connectivity blueprints derived using the macaque data.

**Figure 6.**
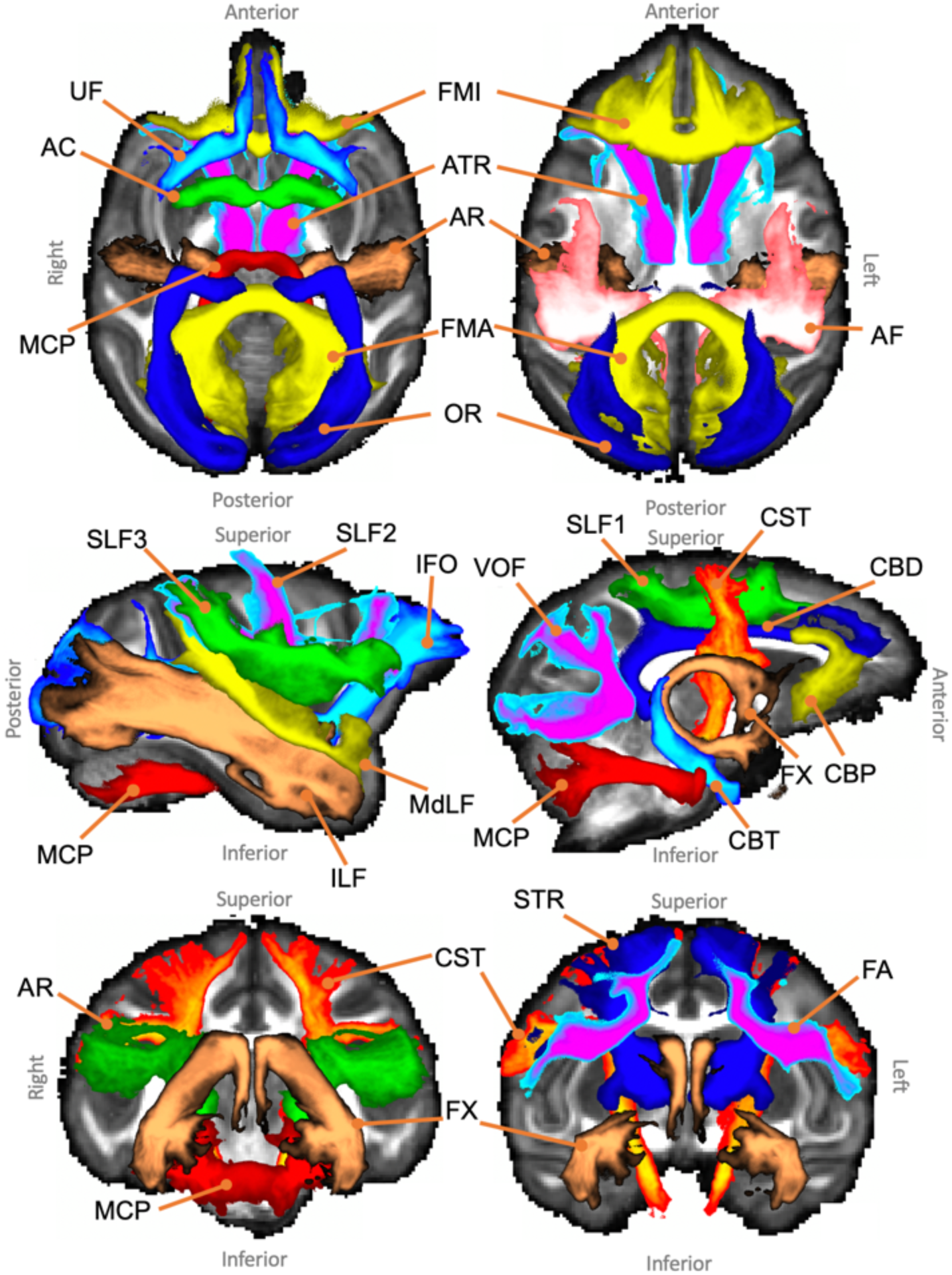
Axial, sagittal and coronal maximal intensity projections of the population percentage tract atlases for the macaque subjects (varying maximal intensity projection window lengths are applied to different tracts for visualisation purposes, display range = 30%-100% of population coverage). Association fibre bundles: Arcuate Fasciculus (AF), Frontal Aslant Tract (FA), Inferior Longitudinal Fasciculus (ILF), Inferior Fronto-Occipital Fasciculus (IFO), Middle Longitudinal Fasciculus (MdLF), Superior Longitudinal Fasciculus I, II and III (SLF), Uncinate Fasciculus (UF) and Vertical Occipital Fasciculus (VOF). Projection fibre bundles: Acoustic Radiation (AR), Anterior Thalamic Radiation (ATR), Corticospinal Tract (CST), Optic Radiation (OR) and Superior Thalamic Radiation (STR). Limbic fibre bundles: Cingulum Bundle: Peri-genual (CBP), Cingulum Bundle: Temporal (CBT), Cingulum Bundle: Dorsal (CBD) and Fornix (FX). Commissural fibre bundles: Anterior Commissure (AC), Forceps Major (FMA) and Forceps Minor (FMI). Tract atlases are created by averaging binarised (threshold of 0.1%) normalised tract density maps across subjects.

**Figure 7.**
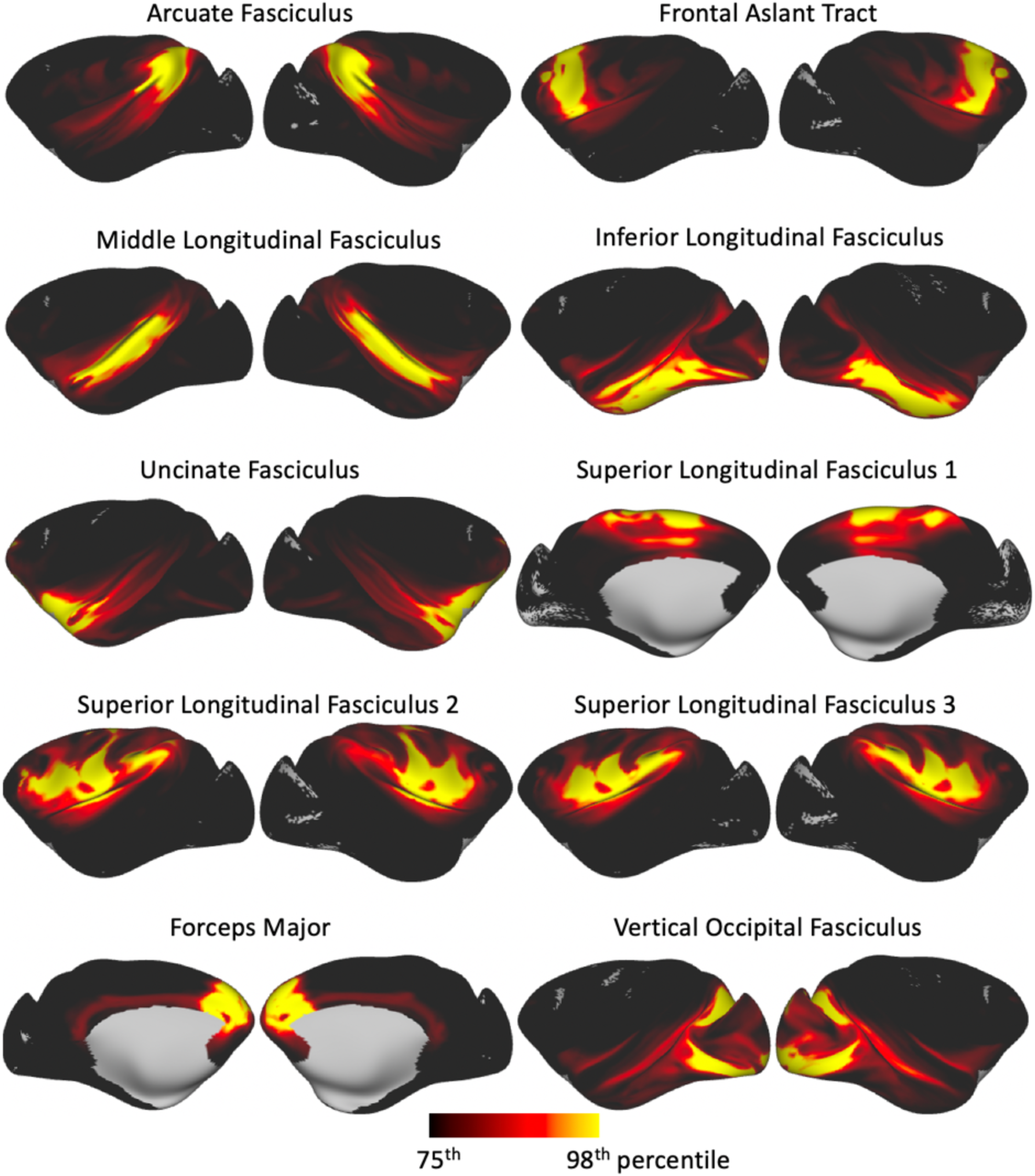
White-grey matter boundary endpoints for a subset of tracts (i.e. columns of the average connectivity blueprint) derived from the macaque subjects.

### Tract lateralisation

Figure 8 illustrates tract-lateralisation estimates for a subset of tracts in the human brain, for which lateralisation has been previously reported in the literature. As shown in Figure 8, the AF is left-lateralised and IFO, MdLF and SLF3 are right-lateralised in both the HCP and the UK Biobank cohorts. SLF1 is symmetric in the HCP cohort but reaches rightward significance in the UK Biobank cohort. SLF2 is also variable across cohorts with left-lateralisation in the HCP and right-lateralisation in the UK Biobank.

**Figure 8.**
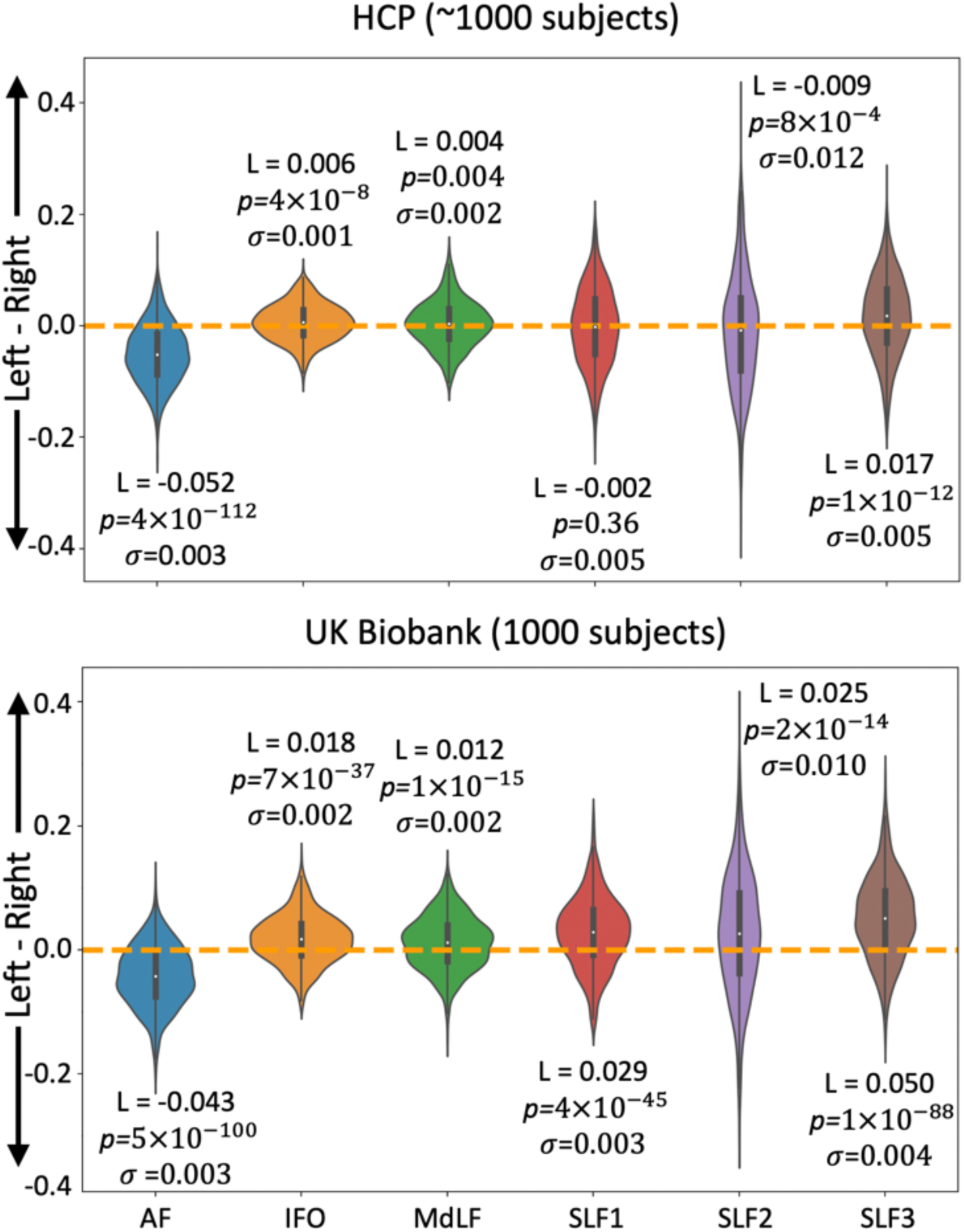
Summary of WM tract lateralisation for the arcuate fasciculus (AF), inferior fronto-occipital fasciculus (IFO), middle longitudinal fasciculus (MdLF) and the superior longitudinal fasciculi (SLFs) using the HCP (top) and UK Biobank (bottom) data. L is the cohort median WM tract lateralisation, p is the p-value obtained from the Mann-Whitney *U* test and σ is the variance for the given WM tract lateralisation. A threshold value of 0.5% has been used to binarise tracts and obtain their volume. Corrected p-value is 0.05/12 = 0.0042.

In addition to volume-based measures of lateralisation, inter-hemispheric differences on connectivity patterns were also assessed on the white-grey matter boundary surface using the tracts-derived connectivity blueprints. In an approach similar to (Mars et al., 2018b), Kullback-Leibler (KL) divergence was calculated to explore connectivity similarity between the two hemispheres of the human brain. For every location on the right hemisphere surface, the minimum KL divergence value assesses the most similar connectivity pattern on the left hemisphere. In doing so, we can probe cortical locations that demonstrate dissimilar connection patterns between left and right hemispheres and assess which tracts are contributing to these dissimilarities. In areas of high minimum KL divergence, we would expect to observe differences in the tract contribution profiles between the corresponding vertices. Figure 9a shows the minimum KL divergence values obtained for all white-grey matter boundary surface locations, overlaid with a subset of the Glasser parcellation (Glasser et al., 2016). Regions of high left-right dissimilarity in connectivity patterns were generally confined to frontal and temporo-parietal junction (TPJ) regions. Figure 9b shows examples of the tract contribution to the connection pattern of specific white-grey matter boundary locations on the right hemisphere and how these compare with the connection patterns of the best matching location on the left hemisphere. Three examples are shown corresponding to varying degrees of inter-hemispheric dissimilarity. It can be seen that high dissimilarity was mediated by tracts that were found to be lateralised. For example, inter-hemispheric differences for a selected voxel in IFSa were primarily driven by differences in how the lateralised SLF3 contributes to its connectivity pattern. For a mid-range dissimilarity vertex, selected in the temporo-parietal-occipital junction (TPOJ1), we can see that small inter-hemispheric divergence was driven primarily by the AF and SLF3. Conversely, regions of low dissimilarity, such as the fourth visual area (V4), show little inter-hemispheric difference in the tracts contributing to the connectivity profile.

**Figure 9.**
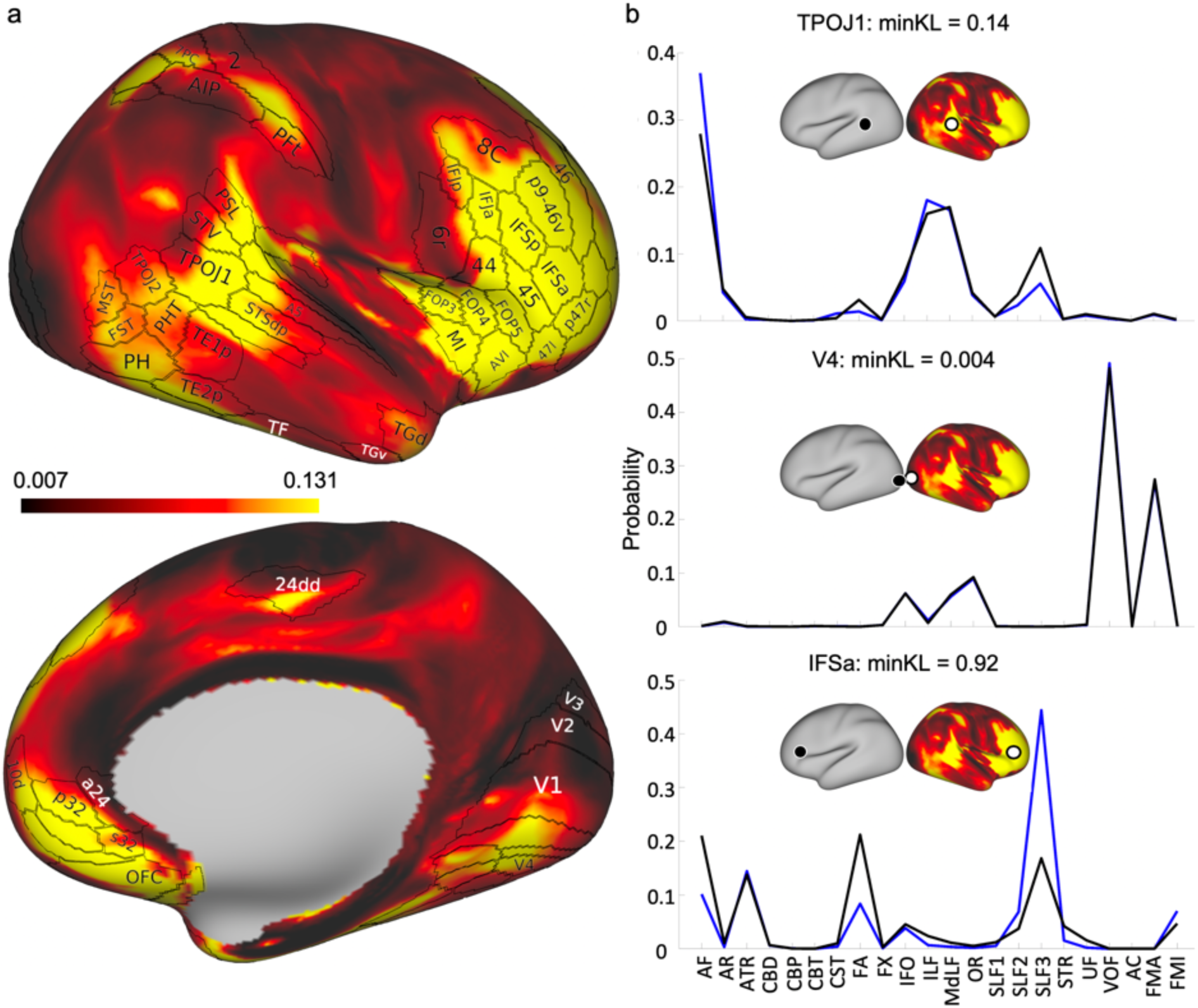
The minimum Kullback-Leibler (KL) divergence between the right and left hemispheres. (a) The minimum KL divergence on the right hemisphere with a subset of the Glasser parcellation (Glasser et al., Nature 2016) highlighting regions of dissimilarity. (b) By selecting vertices of interest from the right hemisphere (white circle) in (a) and extracting the vertex in the left hemisphere (black circle) with the greatest similarity, it is possible to investigate how differences in tract contribution to location connectivity contribute to divergence. Black lines correspond to the tract contributions to the vertex on the right hemisphere and blue lines for the left hemisphere. For example, in V4 (middle), the minimum KL-divergence is small which is reflected by the almost identical underlying tract contributions. Regions with mid-to high-range dissimilarity – TPOJ1 (top) and IFSa (bottom) – are seen to have greater differences in their underlying tract contributions, primarily driven by differences in AF and SLFs.

### Twinship-induced similarities

To explore whether the tractography protocols preserved individual variability in WM anatomy, we compared tract reconstructions for 72 pairs of monozygotic twins, dizygotic twins, non-twin siblings and unrelated subjects from the HCP. Figure 10 shows the distributions of the average subject-wise cross correlations (i.e. average across tracts for each subject) for each group. As shown in Figure 10, monozygotic twin pairs, on average (median), have a higher correlation (0.588, standard deviation = 0.036) with their corresponding twin compared to dizygotic twin pairs (0.545, standard deviation = 0.031), non-twin sibling pairs (0.543, standard deviation = 0.034), and unrelated subject pairs (0.507, standard deviation = 0.029). A Kruskal-Wallis test demonstrates statistically significant differences between subgroup medians: χ^2^ = 122.3, p = 2×10^-26.

**Figure 10.**
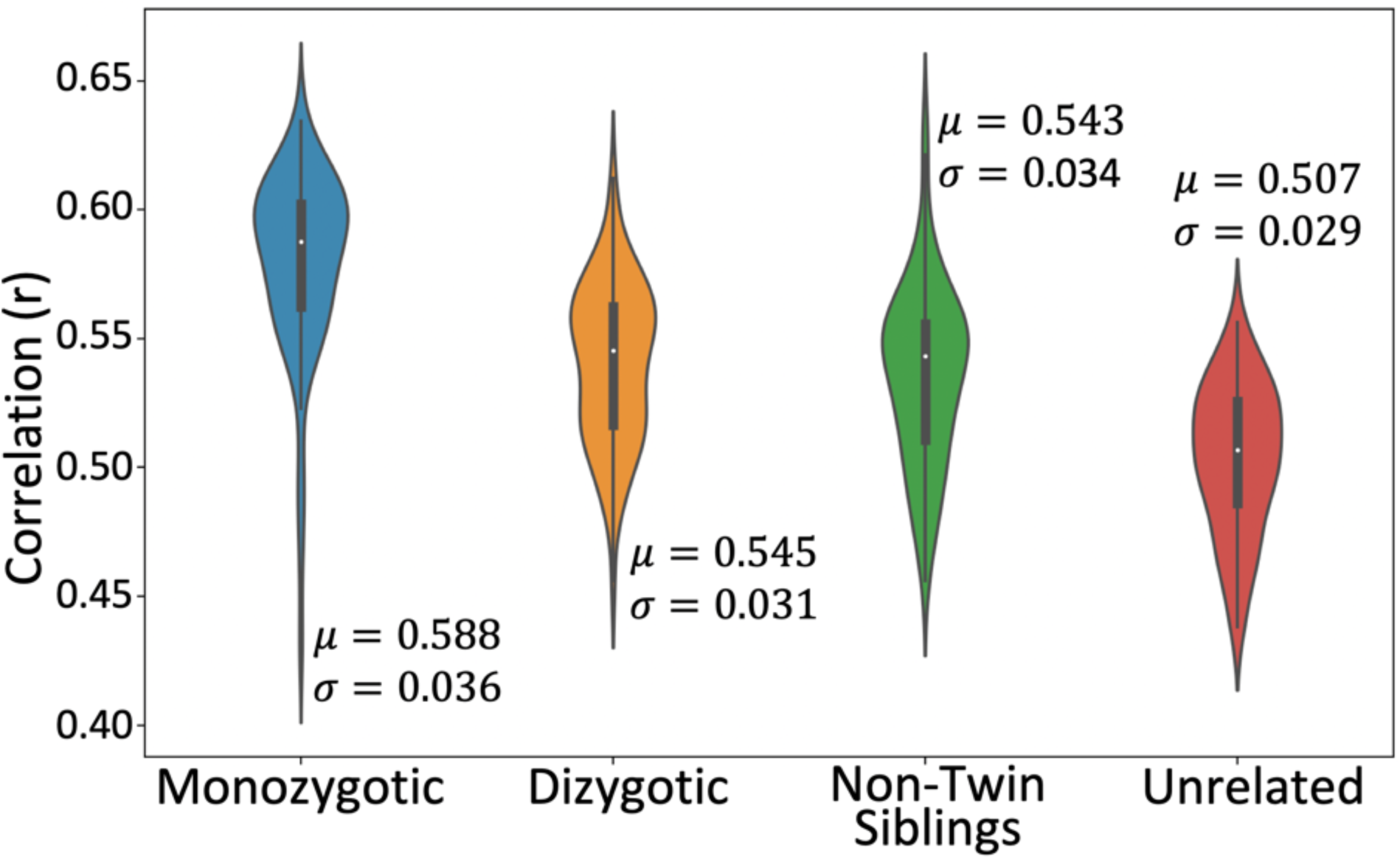
Twin/non-twin WM tract similarity using 72 subject pairs per group. Correlations are performed on normalised tract density maps with a threshold of 0.5%. μ is the group median across tracts and subjects and σ is the standard deviation. A Kruskal-Wallis test is used to determine whether the groups come from the same median: χ^2^ = 122.3, p = 2×10^-26, mean ranks = 222.6 (monozygotic), 145.1 (dizygotic), 134.4 (non-twin siblings) and 69.9 (unrelated).

### Respecting atypical anatomy

In addition to exploring the similarity of tract reconstructions in twins, we further investigated whether the automated tractography respected individual variability in the case of anatomical abnormalities. A subset of subjects with gross anatomical abnormalities were identified using the HCP quality control. Figure 11 and Supplementary Figure 5 give examples of these subjects and highlight the differences between the cohort-averaged tracts and the individual subject tractography results, which reflect the presence of cavernomas, cysts and developmental venous anomalies (DVAs) in WM. Individual tractography results respect the anatomical abnormalities, while atlas-based tracts mask them.

**Figure 11.**
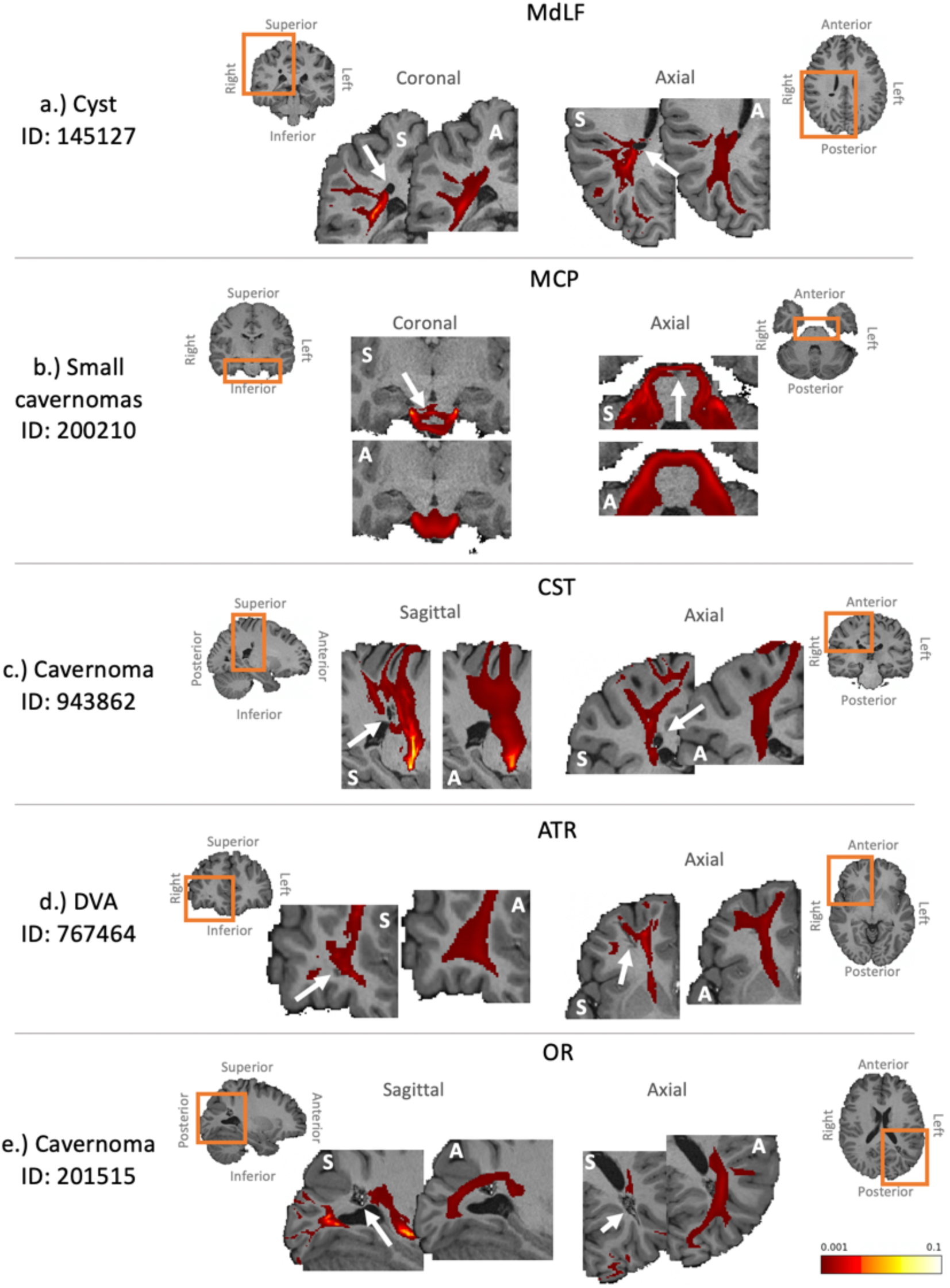
Examples of tractography results for a subset of the subjects found to have anatomical abnormalities, demonstrating that cohort-averaged tracts do not respect underlying anatomical abnormalities. In each case the anatomical abnormality (as described on the HCP quality control website) is given and indicated by the white arrows, and affected tracts are presented. Both the individual subject’s extracted tract (indicated by “S”) and the corresponding cohort-averaged tract (indicated by “A”) are overlaid on a zoomed in individual T1-weighted scan (the orange boxes show the zoomed regions). Tracts are displayed with a threshold of 0.1%. a.) a cyst in the right parietal lobe affecting the MdLF. b.) small cavernoma in the brain stem affecting the MCP. c.) a small cavernoma in the right parietal lobe affecting the CST. d.) a developmental venous anomaly (DVA) in the right frontal lobe affecting the ATR. e.) a cavernoma in the left occipital lobe affecting the OR.

## Discussion

We have presented a new toolbox (XTRACT) for automated probabilistic tractography along with standardised protocols for extracting white matter bundles in the human and the macaque brain. We have demonstrated that the protocols are robust when applied to data of varying image quality and to data from a non-human primate species. We have generated human WM tract atlases using an order of magnitude more data than previous efforts, as well as macaque atlases using a small number of, however high-quality ex vivo, datasets. We have performed indirect validation illustrating that reconstructed tracts are left/right asymmetric, when they are expected to be, based on previous literature. We have also shown that despite automatically generating tracts using standard-space protocols, the protocols respect the underlying individual variability, as reflected in twinship-induced similarities and in respecting anatomical abnormalities. The toolbox, tractography protocols and atlases are freely and openly available as a part of FMRIB’s software library (FSL) (version 6.0.2 and later).

A current issue in the field of tractography is that protocol definitions in the literature often lack detail or are designed without data-sharing in mind. Here, we offer a platform for direct sharing of standardised protocol masks and tract atlases, allowing for the standardisation of tractography protocols across studies and aiding reproducibility. Moreover, we made the protocol definitions generalisable across species to directly facilitate comparative anatomy studies. Anatomists/tractographers may exchange their tract definitions/methods and hopefully converge on consensus protocols, as is the focus of a current multi-centre consortium on defining and standardising WM tractography (cf. Schilling, Landman, Descoteaux et al.). We believe that XTRACT can contribute to these efforts.

A platform for tractography protocol definitions is also presented in (Wassermann et al., 2016), where a white matter query language (WMQL) is devised. Our approach uses a similar logic to WMQL, in the sense that it also relies on masks and Boolean operations on streamlines going through those masks. However, our protocols are generalisable and their utility in both the human and non-human primate brain has been demonstrated here. Also, a main conceptual difference is that we don’t rely on an automated grey matter (GM) parcellation, such as WMQL, but instead on hand-defined white matter masks. WMQL relies on seeding in WM and defining cortical endpoints through subject-wise brain parcellations. Using this approach for tracking into GM has challenges; bottlenecks exist in the cortical gyrus, after which fibres fan out, and most tractography algorithms have issues resolving this fanning (Maier-Hein et al., 2017). To mitigate these issues, our protocols are not dependent on GM masks. We instead focus on reconstructing the main body of the tracts of interest using ROIs mostly in WM. In order to obtain cortical termination points of WM tracts, we take the opposite approach of tracking from the GM surface towards the WM, thus following the direction in which the fibres are expected to merge, rather than to fan out. We then multiply the surface-to-WM tractrogram with that of the body of the tract to create the WM by GM surface projections of the tract (connectivity blueprint). This avoids some of the major problems associated with tracking towards the surface (Eichert et al., 2019b; Mars et al., 2018b).

We reconstructed tracts using imaging datasets of different quality and we generated atlases for both the Human Connectome Project (HCP) and the UK Biobank cohorts. Comparisons of tract reconstructions within and across the human cohorts demonstrate that the method and protocols are robust across subjects and against data quality. The HCP and UK Biobank cohorts provide examples of high-quality data and more typical quality data respectively. Within cohort comparisons reveal similar inter-subject tract correlations across the varying quality data, with greater inter-subject tract correlations observed in the UK Biobank. This may reflect a reduced level of detail in the lower resolution UK Biobank data compared to the HCP data, but also differences in the mean age of subjects in the two cohorts. In addition, we have generated atlases using a smaller cohort of macaques. To compensate for the small number of subjects, we used high-quality and high-resolution ex vivo data. The respective results demonstrate the generalisability of our method to the macaque brain. Recent efforts to obtain macaque data from larger cohorts (HCP-style protocols) are ongoing (Autio et al., 2019; Milham et al., 2018) and our tools will be a useful resource for these new initiatives for the non-human primate brain (Thiebaut de Schotten et al., 2019).

As a means to indirectly validate our results, we investigated left-right tract lateralisation. We compared our lateralisation results to *a priori* knowledge from the literature. For both human cohorts (HCP and UK Biobank), we found that reconstructed AF is strongly left-lateralised, while SLF3, IFO and MDLF were right-lateralised, as expected from the literature (Eichert et al., 2019b; Hau et al., 2016; Hecht et al., 2015; Nowell et al., 2016; Panesar et al., 2018; Thiebaut de Schotten et al., 2011a). Results were less clear-cut for SLF1 and SLF2, where prior studies (with much fewer numbers of subjects) are inconclusive (Hecht et al., 2015; Howells et al., 2018; Thiebaut de Schotten et al., 2011a). This may be due to the large variance observed (in the case of SLF2), perhaps reflecting some underlying interaction, such as handedness (Howells et al., 2018).

We performed further sanity checks by investigating lateralisation using the connectivity blueprint that we obtained from the reconstructed tracts. By using the KL divergence between connectivity patterns to assess inter-hemispheric dissimilarity, we identified that regions associated with language, known to be lateralised (Hiscock and Kinsbourne, 2008), have dissimilar connectivity patterns across the two hemispheres (Figure 9a).

Whilst being a robust automated method for the consistent reconstruction of tracts, our method also respected the underlying anatomical variation. We demonstrated this by assessing inter-subject tract similarity in monozygotic twins, dizygotic twins, non-twin siblings and unrelated subject pairs. Our results show greater similarity in twin pairs compared to unrelated pairs, as would be expected from the heritability literature (Bohlken et al., 2014; Shen et al., 2014).

We further demonstrated that the automated method respects underlying anatomical variation by exploring how tractography results differ from the cohort-averaged results in the case of subjects with anatomical abnormalities. In the presented cases, we show that the toolbox is capable of respecting atypical anatomy. These example cases are of course not exhaustive but offer insight into how the toolbox performs in the presence of relatively small anatomical abnormalities, induced by pathology. We should point out however, that our approach is registration-based, therefore tracking performance in the presence of geometrical malformations is likely to depend on the extent and location of the abnormality and the influence it has to registration (Zhang et al., 2008). Smaller malformations, such as focal WM abnormalities/hyperintensities, are less likely to reduce reliability, particularly given the use of inclusive ROIs in our protocol definitions, as demonstrated in Figure 11 and in agreement with similar findings in recent studies that used tract-derived features (Horbruegger et al., 2019; Ressel et al., 2018). For larger malformations (such as tumours, oedema), even if some compensation can be achieved by performing conditional registration (i.e. by masking out large malformations in the computation of the warp fields), reductions in tracking accuracy may occur and case-specific alternatives may need to be considered (Fekonja et al., 2019). Nevertheless, a range of pathologies do not induce geometrical malformations (for instance the spectrum of psychiatric/neurodevelopmental/mental health disorders) and we expect our approach for delineating major WM tracts to be robust in such cases.

## Conclusions

In conclusion, we have developed and demonstrated a set of robust and standardised tractography protocols for cross-species automated delineation of white matter bundles, along with a platform to use them. The demonstrated toolbox (XTRACT) is freely available along with the tractography protocols and human/macaque tract atlases as a part of FMRIB’s software library (FSL version 6.0.2 and later). Given the benefits with regards to data and protocol sharing, we expect that this toolbox will aid reproducibility in the field of tractography and facilitate comparative neuroanatomy studies.

## Acknowledgements

S.W. is supported by a Medical Research Council PhD Studentship (MR/N013913/1). K.L.B. was supported by a Marie Skłodowska-Curie Individual Fellowship Grant MSCA-IF [750026]. R.B.M. is supported by the Biotechnology and Biological Sciences Research Council (BBSRC) UK [BB/N019814/1] and the Netherlands Organization for Scientific Research NWO [452-13-015]. J.S. was supported by a Sir Henry Dale Wellcome Trust Fellowship (105651/Z/14/Z). G.D. is supported by an MRC Career Development Fellowship (MR/K006673/1). The work was also supported by grant EP/L023067/1 from the UK Engineering and Physical Sciences Research Council (EPSRC) and by a Wellcome Trust grant [217266/Z/19/Z]. Human datasets were provided in part by a) The Human Connectome Project, WU-Minn Consortium (Principal Investigators: David Van Essen and Kamil Ugurbil; 1U54MH091657) funded by the 16 NIH Institutes and Centers that support the NIH Blueprint for Neuroscience Research; and by the McDonnell Center for Systems Neuroscience at Washington University and b) The UK Biobank Resource under Application Number 43822. The computations described in this paper were performed using the University of Nottingham’s Augusta HPC service and the Precision Imaging Beacon Cluster, which provide High Performance Computing service to the University’s research community. The Wellcome Centre for Integrative Neuroimaging is supported by core funding from the Wellcome Trust [203139/Z/16/Z].

## Code and data availability statement

All the data and code used in this work are (or will soon be) publicly available.

Regarding the datasets, there are three sources utilised in this study: a) The WU-Minn Human Connectome Project data (HCP) (available via https://www.humanconnectome.org/), the UK Biobank (available via https://www.ukbiobank.ac.uk/), and locally acquired ex vivo macaques datasets. The macaque datasets will be made available via the PRIME-DE database (http://fcon_1000.projects.nitrc.org/indi/indiPRIME.html).

The XTRACT software toolbox and standardised tractography protocols are already freely available via FMRIB’s software library (FSL) (version 6.0.2 and later: https://fsl.fmrib.ox.ac.uk/fsl/fslwiki/XTRACT) along with a tool for viewing the tractography results (XTRACT_viewer). Atlases (human and macaque WM tracts and connectivity blueprints) generated as a part of this work are currently available via GitHub (https://github.com/SPMIC-UoN/XTRACT_atlases) and will be made available in the next public release of FSL. Code used to perform the Kullback-Leibler divergence analysis is adapted from our previous work and can be found here: https://git.fmrib.ox.ac.uk/rmars/comparing-connectivity-blueprints.

## Footnotes

1 XTRACT toolbox and human/macaque tractography protocols are available in FSL version 6.0.2 and later (www.fmrib.ox.ac.uk/fsl/XTRACT). Tract atlases will be made available in the next release of FSL and are currently available via GitHub (https://github.com/SPMIC-UoN/XTRACT_atlases).

2 HCP quality control website - https://wiki.humanconnectome.org/pages/viewpage.action?pageId=88901591

3 PRIMatE Data Exchange - http://fcon_1000.projects.nitrc.org/indi/indiPRIME.html

4 Macaque registrations to the F99 atlas are performed using a custom configuration file. The F99 atlas volumes and surfaces are distributed within XTRACT, along with the custom configuration file.

