## Supplementary Material for "XTRACT - Standardised protocols for automated tractography in the human and macaque brain"

| Tract | Wakana[1] (N=4) | Catani[2] (N=12) | Hua[3] (N=28) | Zhang[4] (N=10) | Thiebaut de Schotten[5] (N=40) | Yendiki[6] (N=67) | de Groot[7] (N=60) | Wassermann[8] (N=97) |
|---|---|---|---|---|---|---|---|---|
| Acoustic Radiation | | | | | | | ✓ | |
| Anterior Commissure | | ✓ | | | ✓ | | | |
| Anterior Thalamic Radiation | ✓ | | ✓ | ✓ | | ✓ | ✓ | |
| Arcuate Fasciculus | | ✓ | | | ✓ (Arcuate and anterior, long and posterior segments) | | | ✓ |
| Cingulum subsection: Dorsal | ✓ | | ✓ | ✓ | | ✓ | ✓ | |
| Cingulum subsection: Peri-genual | ✓ | ✓ | ✓ | ✓ | ✓ | ✓ | ✓ | ✓ |
| Cingulum subsection: Temporal | | | | | | | | |
| Corpus Collosum | | ✓ | | | ✓ | | | ✓ (Rostrum, rostral body, anterior midbody, posterior midbody, isthmus) |
| Corpus Collosum: Forceps Major | ✓ | | ✓ | ✓ | | ✓ | ✓ | ✓ |
| Corpus Collosum: Forceps Minor | ✓ | | ✓ | ✓ | | ✓ | ✓ | ✓ |
| Cortico-Ponto-Cerebellar | | | | | ✓ | | | |
| Corticospinal Tract | ✓ | | ✓ | ✓ | ✓ | ✓ | ✓ | ✓ |
| Extreme Capsule | | | | | | | | ✓ |
| Fornix | | ✓ | | | ✓ | | | |
| Inferior Cerebellar Peduncle | | ✓ | | | | | | |
| Inferior Fronto-Occipital Fasciculus | ✓ | ✓ | ✓ | ✓ | ✓ | | ✓ | ✓ |
| Inferior Longitudinal Fasciculus | ✓ | ✓ | ✓ | ✓ | ✓ | ✓ | ✓ | ✓ |
| Internal Capsule/Corona Radiata | | ✓ | | | ✓ | | | |
| Medial Lemniscus | | | | | | | ✓ | |
| Middle Cerebellar Peduncle | | ✓ | | | | | ✓ | |
| Middle Longitudinal Fasciculus | | | | | | | | ✓ |
| Optic Radiation/Posterior Thalamic Radiation | | | | | ✓ | | ✓ | ✓ |
| Spino-Cerebellar Tract | | | | | ✓ | | | |
| Striato-Fronto-Orbital | | | | | | | | ✓ |
| Striato-Prefrontal | | | | | | | | ✓ |
| Striato-Premotor | | | | | | | | ✓ |
| Striato-Precentral | | | | | | | | ✓ |
| Striato-Postcentral | | | | | | | | ✓ |
| Striato-Parietal | | | | | | | | ✓ |
| Striato-Occipital | | | | | | | | ✓ |
| Superior Cerebellar Peduncle | | ✓ | | | ✓ | | | |
| Superior Longitudinal Fasciculus I | ✓ | | ✓ | ✓ | | ✓ | ✓ | ✓ |

| | | | | | | | | |
|---|---|---|---|---|---|---|---|---|
| Superior Longitudinal Fasciculus II | (SLF and the temporal component of the SLF) | | (SLF and the temporal component of the SLF) | (SLF and the temporal component of the SLF) | | (SLF and the temporal component of the SLF) | | ✓ |
| Superior Longitudinal Fasciculus III | | | | | | | | ✓ |
| Superior Thalamic Radiation | | | | | | | ✓ | |
| Thalamo-Prefrontal | | | | | | | | ✓ |
| Thalamo-Premotor | | | | | | | | ✓ |
| Thalamo-Precentral | | | | | | | | ✓ |
| Thalamo-Postcentral | | | | | | | | ✓ |
| Thalamo-Parietal | | | | | | | | ✓ |
| Thalamo-Occipital | | | | | | | | ✓ |
| Uncinate Fasciculus | ✓ | ✓ | ✓ | ✓ | ✓ | ✓ | ✓ | ✓ |
| **Total:** | 20 | 19 | 22 | 22 | 31 | 18 | 27 | 57 |

**Supplementary Table *1*.** A brief review of the protocols previously defined in the literature. N=number of subjects used.

[1]Wakana, S., Jiang, H., Nagae-Poetscher, L.M., van Zijl, P.C., Mori, S., 2004. Fiber Tract–based Atlas of Human White Matter Anatomy. Radiology.
[2]Catani, M., Thiebaut de Schotten, M., 2008. A diffusion tensor imaging tractography atlas for virtual in vivo dissections. Cortex.
[3]Hua, K., Zhang, J., Wakana, S., Jiang, H., Li, X., Reich, D.S., Calabresi, P.A., Paker, J.J., van Zijl, P.C.M., Mori, S., 2008. Tract probability maps in stereotaxic spaces: Analysis of white matter anatomy and tract-specific quantification. NeuroImage.
[4]Zhang, W., Olivi, A., Hertij, S.J, van Zijl, P., Mori, S., 2008. Automated fiber tracking of human brain white matter using diffusion imaging. NeuroImage.
[5]Thiebaut de Schotten, M., Ffytche, D.H., Bizzi, A., Dell'Acqua, F., Allin, M., Walshe, M., Murray, R., Williams, S.C., Murphy, D.G., Catani, M., 2011. Atlasing location, asymmetry and inter-subject variability of white matter tracts in the human brain with MR diffusion tractography. NeuroImage.
[6]Yendiki, A., Panneck, P., Srinivasan, P., Stevens, A., Zöllei, L., Augustinack, J., Wang, R., Salat, D., Ehrlich, S., Behrens, T., Jbabdi, S., Gollub, R., Fischl, B., 2011. Automated probabilistic reconstruction of white-matter pathways in health and disease using an atlas of the underlying anatomy. Front. Neuroinform.
[7]de Groot, M., Vernooij, M.W., Klein, S., Ikram, M.A., Vos, F.M., Smith, S.M., Niessen, W.J., Andersson, J.L., 2013. Improving alignment in Tract-based spatial statistics: Evaluation and optimization of image registration. NeuroImage.
[8]Wassermann, D., Makris, N., Rathi, Y., Shenton, M., Kikinis, R., Kubicki, M., Westin, C.F., 2016. The white matter query language: a novel approach for describing human white matter anatomy. Brain Struct. Funct.

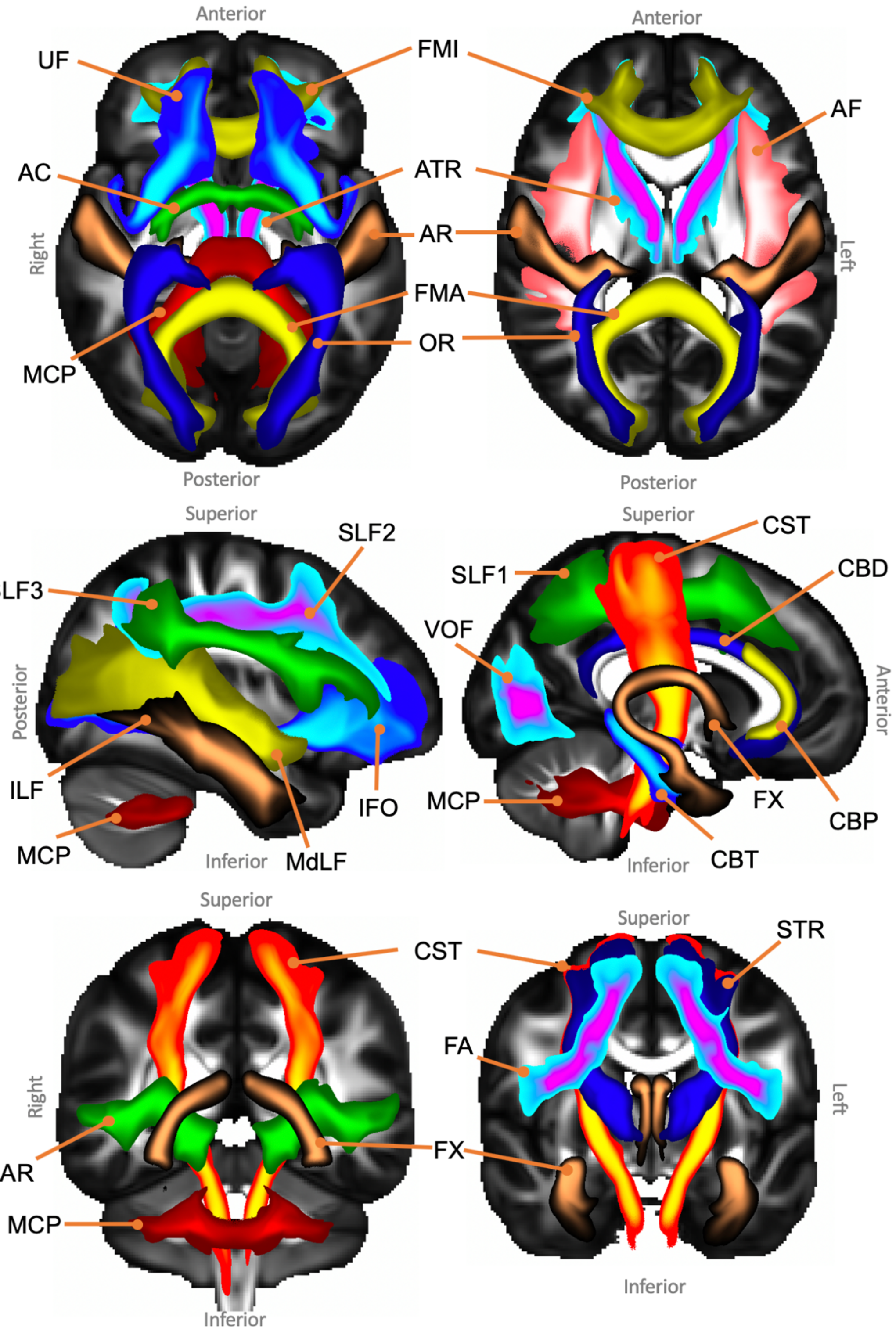


**Supplementary Figure 1.** Axial, sagittal and coronal maximal intensity projections of the population percentage tract atlases for the UK Biobank subset (varying maximal intensity projection window lengths are applied to different tracts for visualisation purposes, display range = 5%-100% of population coverage). Association fibre bundles: Arcuate Fasciculus (AF), Frontal Aslant Tract (FA), Inferior Longitudinal Fasciculus (ILF), Inferior Fronto-Occipital Fasciculus (IFO), Middle Longitudinal Fasciculus (MdLF), Superior Longitudinal Fasciculus I, II and III (SLF), Uncinate Fasciculus (UF) and Vertical Occipital Fasciculus (VOF). Projection fibre bundles: Acoustic Radiation (AR), Anterior Thalamic Radiation (ATR), Corticospinal Tract (CST), Optic Radiation (OR) and Superior Thalamic Radiation (STR). Limbic fibre bundles: Cingulum Bundle: Peri-genual (CBP), Cingulum Bundle: Temporal (CBT), Cingulum Bundle: Dorsal (CBD) and Fornix (FX). Commissural fibre bundles: Anterior Commissure (AC), Forceps Major (FMA) and Forceps Minor (FMI). Tract atlases are created by averaging binarised (threshold of 0.1%) normalised tract density maps across subjects.

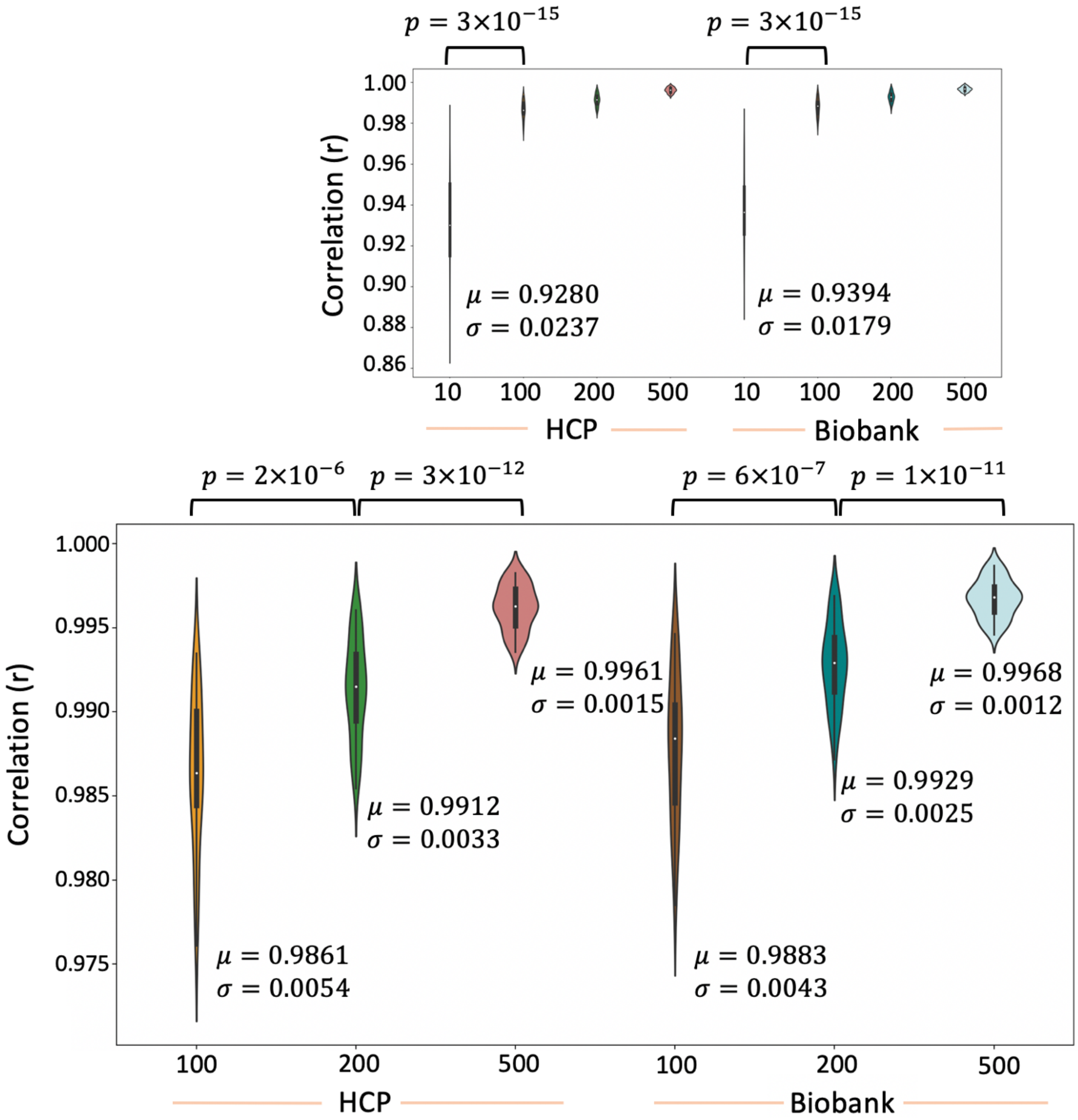


**Supplementary Figure 2.** Summary of the tract-wise correlations of varying sample-size atlases to 1000-subject atlas for the HCP and the UK Biobank datasets. Each atlas is created by binarising (threshold of 0.1%) each of the subject's normalised tract density maps and then averaging across subjects. The full range of sample sizes used (10, 100, 200 and 500) is shown at the top and a zoomed-in version highlighting the differences between the larger sample sizes (100, 200 and 500) is shown at the bottom. $\mu$ is the group mean across subjects and $\sigma$ is the standard deviation. Significance is obtained via Mann-Whitney U test. Corrected p-value is 0.05/6 = 0.0083.

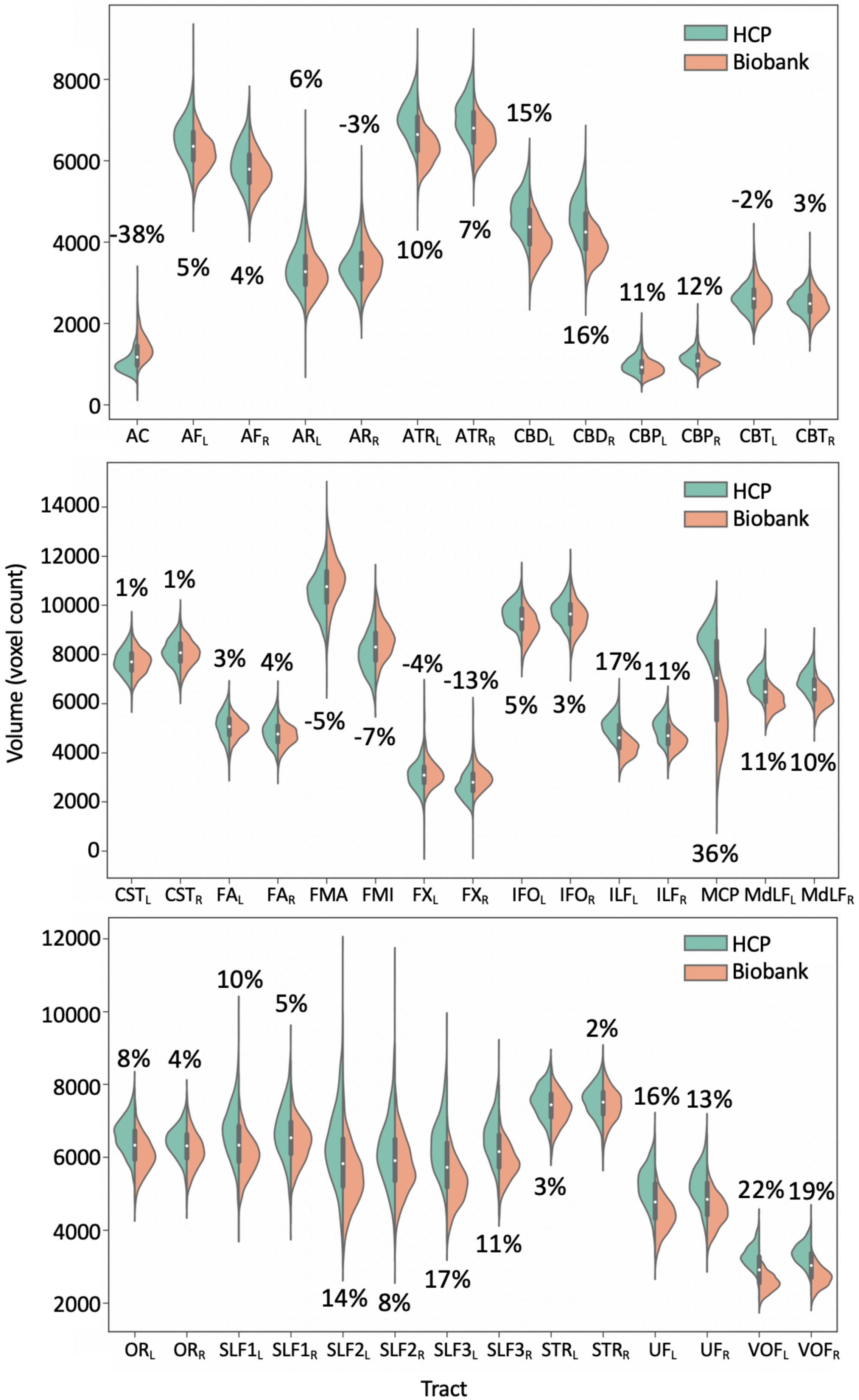


**Supplementary Figure 3.** Comparisons of the cohort-averaged volumes of each tract for the HCP (green) and UK Biobank (orange). Volume is taken as the sum of non-zero voxels following the binarisation of the waytotal normalised tract density maps (threshold of 0.5%). Percentages indicate the percent difference between the average tract volume in the HCP cohort compared to the UK Biobank cohort, relative to the HCP cohort, i.e. $100*((V_{HCP}-V_{Biobank})/V_{HCP})$.

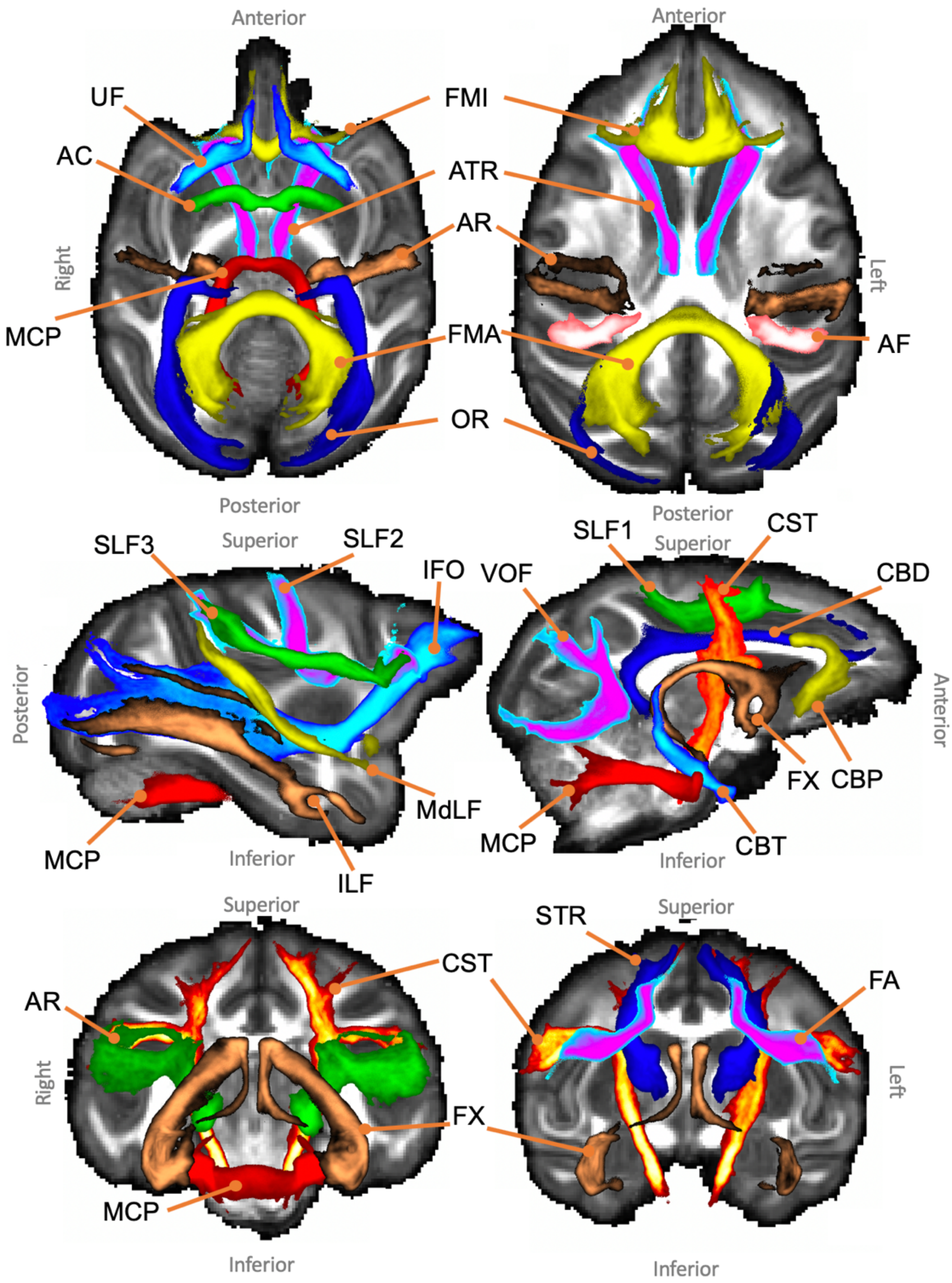


**Supplementary Figure 4.** Axial, sagittal and coronal maximal intensity projections of the population percentage tract atlases for the macaque subjects (varying maximal intensity projection window lengths are applied to different tracts for visualisation purposes, display range = 30%-100% of population coverage). Association fibre bundles: Arcuate Fasciculus (AF), Frontal Aslant Tract (FA), Inferior Longitudinal Fasciculus (ILF), Inferior Fronto-Occipital Fasciculus (IFO), Middle Longitudinal Fasciculus (MdLF), Superior Longitudinal Fasciculus I, II and III (SLF), Uncinate Fasciculus (UF) and Vertical Occipital Fasciculus (VOF). Projection fibre bundles: Acoustic Radiation (AR), Anterior Thalamic Radiation (ATR), Corticospinal Tract (CST), Optic Radiation (OR) and Superior Thalamic Radiation (STR). Limbic fibre bundles: Cingulum Bundle: Peri-genual (CBP), Cingulum Bundle: Temporal (CBT), Cingulum Bundle: Dorsal (CBD) and Fornix (FX). Commissural fibre bundles: Anterior Commissure (AC), Forceps Major (FMA) and Forceps Minor (FMI). Tract atlases are created by averaging binarised (threshold of 0.5%) normalised tract density maps across subjects.

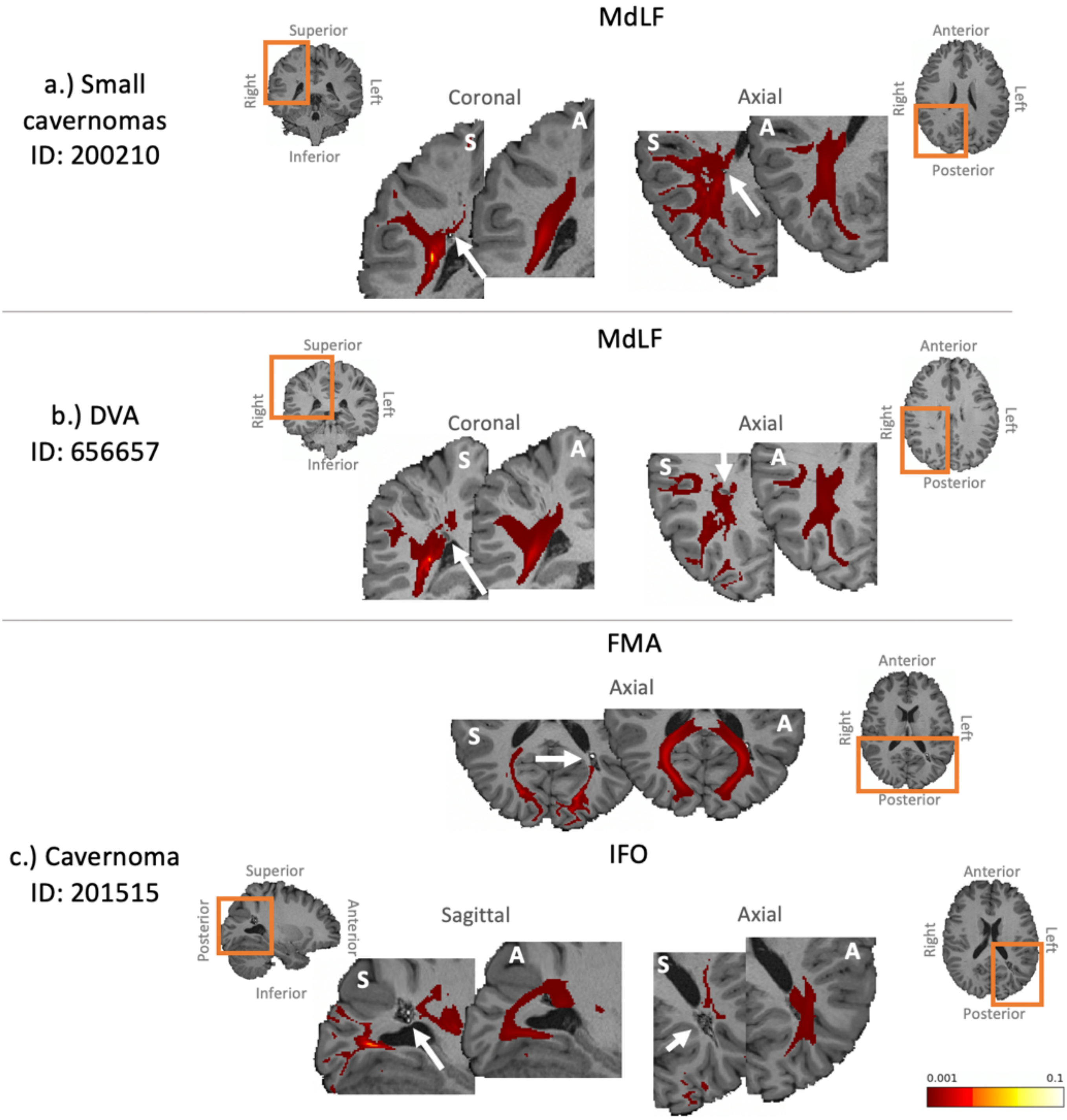


**Supplementary Figure 5.** Further examples of tractography results for a subset of the subjects found to have anatomical abnormalities. a.) small cavernoma in the right parietal lobe (same subject as Figure 11b) affecting the MdLF. b.) a developmental venous anomaly (DVA) in the right parietal lobe affecting the MdLF. c.) a cavernoma in the left occipital lobe (same subject as Figure 11e) affecting the FMA and IFO.
